# Evolutionary Dynamics of G-Quadruplexes in Human and Other Great Ape Telomere-to-Telomere Genomes

**DOI:** 10.1101/2024.11.05.621973

**Authors:** Saswat K. Mohanty, Francesca Chiaromonte, Kateryna D. Makova

**Affiliations:** Molecular, Cellular, and Integrative Biosciences, Huck Institutes of the Life Sciences, Penn State University, University Park, PA 16802, USA; Department of Biology, Penn State University, University Park, PA 16802, USA; Department of Statistics, Penn State University, University Park, PA 16802, USA; Center for Medical Genomics, Penn State University, University Park and Hershey, PA, USA; EMbeDS, Sant’Anna School of Advanced Studies, 56127 Pisa, Italy

## Abstract

G-quadruplexes (G4s) are non-canonical DNA structures that can form at approximately 1% of the human genome. G4s contribute to point mutations and structural variation and thus facilitate genomic instability. They play important roles in regulating replication, transcription, and telomere maintenance, and some of them evolve under purifying selection. Nevertheless, the evolutionary dynamics of G4s has remained underexplored. Here we conducted a comprehensive analysis of predicted G4s (pG4s) in the recently released, telomere-to-telomere (T2T) genomes of human and other great apes—bonobo, chimpanzee, gorilla, Bornean orangutan, and Sumatran orangutan. We annotated tens of thousands of new pG4s in T2T compared to previous ape genome assemblies, including 41,236 in the human genome. Analyzing species alignments, we found approximately one-third of pG4s shared by all apes studied and identified thousands of species– and genus-specific pG4s. pG4s accumulated and diverged at rates consistent with divergence times between the studied species. We observed a significant enrichment and hypomethylation of pG4 shared across species at regulatory regions, including promoters, 5’ and 3’UTRs, and origins of replication, strongly suggesting their formation and functional role in these regions. pG4s shared among great apes displayed lower methylation levels compared to species-specific pG4s, suggesting evolutionary conservation of functional roles of the former. Many species-specific pG4s were located in the repetitive and satellite regions deciphered in the T2T genomes. Our findings illuminate the evolutionary dynamics of G4s, their role in gene regulation, and their potential contribution to species-specific adaptations in great apes, emphasizing the utility of high-resolution T2T genomes in uncovering previously elusive genomic features.

## INTRODUCTION

Most of genomic DNA exists in B form, i.e. a right-handed double helix with 10 base pairs per turn (Watson and Crick 1953). However, certain regions of the genome can form non-B DNA with different structures (Kouzine et al. 2017), which are transient and have lower stability than B DNA. These alternative DNA structures—such as Z-DNA (Mitsui et al. 1970), cruciforms (Panayotatos and Fontaine 1987), H-DNA (Felsenfeld and Rich 1957), bent DNA (Prosseda et al. 2004), and G-quadruplexes (G4s) (Sen and Gilbert 1988)—are involved in various biological processes (reviewed in (Wang and Vasquez 2022)). Among non-B DNA, G4s have gained considerable attention due to their critical role in genomic regulation (Ravichandran et al. 2019; Pavlova et al. 2021; Zhang et al. 2024). Moreover, G4s have emerged as promising therapeutic targets for drug development (Kosiol et al. 2021; Monsen 2023). Both DNA and RNA (Fay et al. 2017) can form G4 structures, which may be either inter-or intramolecular. These structures arise due to formation of Hoogsteen hydrogen bonds (Hoogsteen 1963) among nonconsecutive guanine bases, which assemble into planar G-tetrads. Multiple G-tetrads are stabilized by monovalent cations such as potassium. A G4 structure is comprised of stems and loops: stems consist of runs of three to five consecutive guanines, whereas loops, which can range from 1 to 12 nucleotides, can include other nucleotides (Guédin et al. 2010). In the cell, G4s with bulges in their stems (i.e. G-runs interrupted by non-G nucleotides) can also form (Mukundan and Phan 2013; Papp et al. 2023).

G4s affect genome stability and regulate gene expression (Guiblet, Cremona, et al. 2021; Teng et al. 2021). G4s are abundant at telomeres, and participate in telomere maintenance (Moye et al. 2015; Lin and Yang 2017; Bryan 2020; Xu and Komiyama 2023). G4s have been suggested to be important regulatory elements in DNA replication, albeit in two contrasting ways: (1) G4s are important for replication initiation, and their deletion can impede this process (Valton et al. 2014; Prioleau 2017); (2) G4s can act as barriers to replication progression, leading to replication fork stalling (Stein et al. 2022) and contributing to genomic instability (Sun and Hurley 2010; Lormand et al. 2013). G4s have also been suggested to contribute to genome instability as mutation hotspots during evolution (Guiblet et al. 2018; Guiblet, Cremona, et al. 2021) and in cancer (Georgakopoulos-Soares et al. 2018; Stein and Eckert 2021). Similarly, G4s likely play a dual role in transcription regulation: (1) G4s can serve as binding hubs for transcription factors (Spiegel et al. 2021) and enhance transcription levels (Lee et al. 2020); (2) when present on the template strand, they can stall RNA polymerase, thereby blocking transcription (Broxson et al. 2011; Smestad and Maher 2015). Consistent with their crucial regulatory functions, G4 structures have been shown to evolve under purifying selection at multiple genomic regions (Guiblet, DeGiorgio, et al. 2021).

Dysregulation of G4s has been associated with diseases. These structures have been identified in the promoter regions of several genes linked to cancer, including *MYC* (Madden et al. 2021), *KRAS* (D’Aria et al. 2021), *BCL2* (Del Toro et al. 2009), and *KIT* (Phan et al. 2007). Relatedly, G4s have been used as therapeutic targets for melanoma, leukemia, and pancreatic cancer (Kosiol et al. 2021). Moreover, G4s have been associated with multiple neurodegenerative diseases (reviewed in (Wang et al. 2021) such as X-linked dystonia-parkinsonism (Nicoletto et al. 2024), Alzheimer’s disease (Lammich et al. 2011; Fisette et al. 2012; Crenshaw et al. 2015), Parkinson’s disease (Koukouraki and Doxakis 2016), fragile X syndrome (Zhang et al. 2014; McAninch et al. 2017), Amyotrophic Lateral Sclerosis (Haeusler et al. 2014), and Progressive Myoclonus Epilepsy Type I (Saha and Usdin 2001). Thus, studies of G4s are clinically relevant.

Despite the significance of G4s for mutagenesis and cellular functions, as well as their clinical relevance, their evolution remains understudied. To fill this gap, we conducted a comprehensive analysis of predicted G4s in the recently-released complete, telomere-to-telomere (T2T) genomes of six great apes (Nurk et al. 2022; Rhie et al. 2023; Makova et al. 2024; Yoo et al. 2024)—human (*Homo sapiens*), chimpanzee (*Pan troglodytes*, diverged ∼7 million years ago, MYA, from the human lineage), bonobo (*Pan paniscus*, diverged ∼2.5 MYA from the chimpanzee lineage), gorilla (*Gorilla gorilla*, diverged ∼9 million years ago from the human lineage), Sumatran orangutan (*Pongo abelii*, diverged ∼17 MYA from the human lineage), and Bornean orangutan (*Pongo pygmaeus*, diverged ∼1 MYA from the Sumatran orangutan lineage). The advent of high-quality T2T reference genomes has enabled a detailed analysis of complex and previously inaccessible genomic regions, including telomeres and centromeres. These T2T genomes, assembled with long-read sequencing technologies having low error rates at G4s (Makova and Weissensteiner 2023), can provide accurate sequence information at G4 loci (Rhie et al. 2023; Smeds et al. 2024). Importantly, the use of T2T genomes in great ape research can facilitate the identification of previously unmapped G4s, providing a more complete understanding of how G4s have evolved across species.

In this study, we present a comprehensive analysis of G4s across the T2T genomes of six species of great apes. We established a complete database of predicted G4s, delving into the newly resolved regions of the T2T assemblies, and providing insights into evolutionary patterns of G4 occurrence, conservation, and divergence among great apes. We identified both shared and species-specific pG4s, offering a comparative view of their evolutionary trajectories. Additionally, we examined G4 enrichment across different genomic contexts. Taking advantage of the availability of methylation data, we were able to predict G4 formation in two cell lines. Our findings pave the way for future studies of the evolutionary impact of G4s, their structural dynamics, and their role in ape evolution.

## RESULTS

### Primate T2T assemblies provide a complete catalog of predicted G4 structures, including tens of thousands of new occurrences

We predicted G4 motifs (pG4s) in the T2T genome assemblies of human, bonobo, chimpanzee, gorilla, Bornean orangutan, and Sumatran orangutan (Yoo et al. 2024). We initially evaluated several computational methods proposed to predict G4 motifs (reviewed in (Lombardi and Londoño-Vallejo 2020)) and used a combination of methods that captured both canonical and non-canonical G4 motifs while minimizing false positives and false negatives. In particular, our pG4 discovery pipeline (Fig. 1) utilized both the pqsfinder (Hon et al. 2017) and the G4Hunter (Bedrat et al. 2016) prediction algorithms along with several filtering steps (see Methods for details).

**Figure 1.**
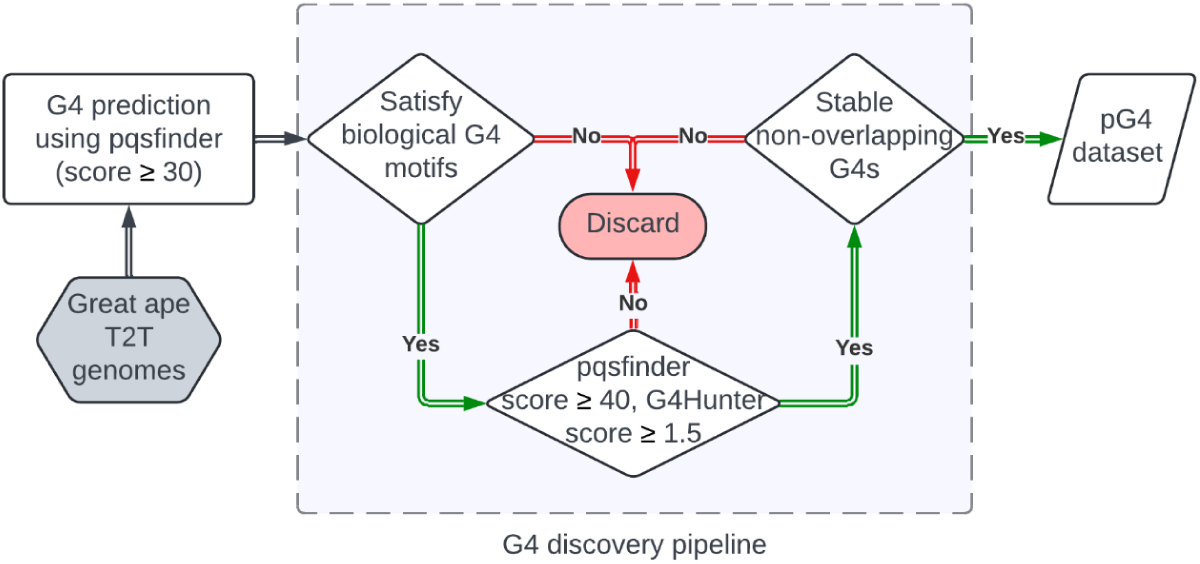
The workflow of the pG4 discovery pipeline. G4s were predicted in each great ape T2T genome using pqsfinder (score threshold ≥ 30). The resulting G4 predictions had to (1) satisfy either standard or bulged G4 motif regular expression, (2) have a minimum score of 40 and 1.5 for pqsfinder and G4Hunter, respectively, and (3) be non-overlapping and have the highest pqsfinder score in comparison with the other pG4s in that region.

We selected the pqsfinder and G4Hunter algorithms because (1) they had the highest accuracy among multiple algorithms tested with an *in vitro* validation dataset (Lombardi and Londoño-Vallejo 2020); and (2) they allowed us to leverage different sequence information. Indeed, to compute a score for a pG4, pqsfinder takes into account the number of tetrads, bulges, and loop lengths (Hon et al. 2017), whereas G4Hunter considers G-richness and G-skewness (Bedrat et al. 2016). To determine the thresholds for each scoring algorithm, we first predicted G4s in the human T2T genome with relaxed parameters, i.e. a pqsfinder score ≥30 and an absolute G4Hunter score ≥0. The distribution of pqsfinder scores exhibited an inflection point at 40 (Fig. S1A), and we considered pG4s with score ≥40 for subsequent analyses (a score of 52 was previously recommended as a minimum threshold (Hon et al. 2017)). The distribution of G4Hunter scores was bell-shaped with no clear inflection point (Fig. S1B). A previous study (Bedrat et al. 2016) reported that G4Hunter precision exceeds 90% above the threshold of 1.5 (for 25-bp G4s), and thus we selected this threshold for our subsequent analysis.

We included pG4s with standard, i.e. [G^3+^L^1−12^]^3+^G^3+^ (Guédin et al. 2010), and bulged, i.e. [GN^0−1^GN^0−1^GL^1−3^]^3+^GN^0−1^GN^0−1^G (Mukundan and Phan 2013), motifs, where L∈{A, T, C, G} and N∈{A, T, C}. We excluded pG4s with uneven motifs, i.e. [G^1−2^N^1−2^]^7+^G^1−2^ (Maity et al. 2020), because such motifs can have two tetrads potentially representing an intermediate G4 form (Zhang et al. 2009); three tetrads were found to be the shortest guanine runs to result in a stable structure formation (Bugaut and Balasubramanian 2008). Additionally, pqsfinder (with a minimum score threshold of 30) could not predict any G4s with uneven motifs.

Using the discovery pipeline described above (Fig. 1), we annotated 769,188–844,654 pG4s in great ape T2T genome assemblies (Table 1), with the lowest number found in human and the highest in gorilla. Among human pG4s, ∼69% were standard and ∼31% were bulged. These proportions were similar for non-human great apes (Table 1). Our final G4 predictions in the human T2T genome contained sequences known to form G4 structures, particularly in the promoter regions of *c-MYC* (Siddiqui-Jain et al.), *TEAD4* (Cozzaglio et al. 2022), *TERT* (Pavlova et al. 2022), *NEIL3* (Fleming et al. 2023), *ZFP42* (Roy et al. 2023) and *SLC6A3* (Nain et al. 2023) genes, and in the body of the *SRC* gene (Rodriguez et al. 2012).

**Table 1.** The number of pG4s annotated in the T2T great ape genomes, with standard and bulged pG4s counted separately, along with the subset of G4s predicted specifically in the newly resolved regions of each species’ T2T genome assembly. Bornean orangutan lacked a previous chromosome-level assembly. (B. orangutan: Bornean orangutan, S. orangutan: Sumatran orangutan)

|  | Standard G4s |  | Bulged G4s |  | G4s in T2T resolved regions | Total G4s |
| --- | --- | --- | --- | --- | --- | --- |
|  | Quantity | Percentage | Quantity | Percentage |  |  |
| **Bonobo** | 552,876 | 69.2% | 245,647 | 30.8% | 76,939 | 798,523 |
| **Chimpanzee** | 547,050 | 69.3% | 241,980 | 30.7% | 64,028 | 789,030 |
| **Human** | 529,843 | 68.9% | 239,345 | 31.1% | 41,236 | 769,188 |
| **Gorilla** | 595,622 | 70.5% | 249,032 | 29.5% | 147,753 | 844,654 |
| **B. orangutan** | 558,169 | 71.1% | 226,921 | 28.9% | – | 785,090 |
| **S. orangutan** | 560,530 | 70.8% | 230,678 | 29.2% | 68,810 | 791,208 |

Importantly, we predicted an additional 41,236 G4s in the newly resolved regions of the human T2T genome assembly (T2T-CHM13v2.0) compared to the previous assembly (GRCh38). The human T2T assembly unveiled a large number of new pG4s at acrocentric chromosomes—13, 14, 15, 21, and 22—and at chromosomes 8 and 10 (Fig. 2). The T2T assemblies for the other non-human great apes, for which previous chromosome-level assemblies were available (all but Bornean orangutan), also unveiled a large number of new pG4s (Fig. S2, Table 1).

**Figure 2.**
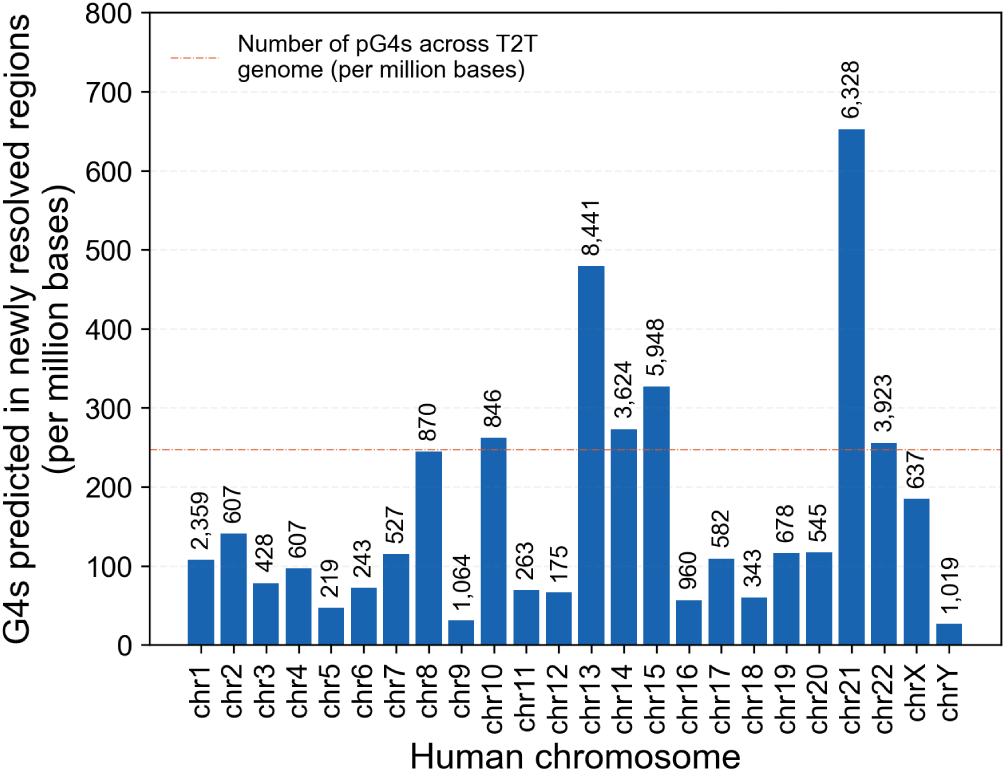
Comparison of predicted G4 density in newly resolved regions of the human T2T genome assembly versus hg38, across all chromosomes. The bars show the number of G4s predicted in the newly resolved regions (per million bases) of the human T2T genome, as compared to the hg38 version, across all chromosomes. The horizontal red line indicates the number of pG4s per million bases across the whole human T2T genome, i.e. the average genome-wide density. The numbers on the top of each bar show the number of pG4s in the newly resolved regions on each chromosome.

### Density of predicted G4s is similar across homologous chromosomes and is positively correlated with gene density

We next examined the density of pG4s across homologous chromosomes in great apes. We constructed a chromosome homology map displaying the homology between great ape and human chromosomes (Fig. 3A). The homology between chromosomes was estimated from whole-genome pairwise alignments (see Methods for details). Structural variants (SVs), such as inversions and duplications, occurring on the same chromosome were not considered. We were able to corroborate several previously known homologous relationships. For instance, human chromosome 9 is homologous to chromosome 11 in *Pan* species (chimpanzee and gorilla) and to chromosome 13 in *Pongo* species (Sumatran and Bornean orangutans) and gorilla. The telomeric fusion of two ancestral acrocentric chromosomes led to the formation of human chromosome 2 (Turleau et al. 1972; Wienberg et al. 1994), which is homologous to chromosomes 12 and 13 in *Pan*, and chromosomes 11 and 12 in *Pongo* and gorilla. In gorilla, a reciprocal translocation occurred between ancestral chromosomes 4 and 19, which are homologous to human chromosomes 5 and 17, respectively (Dutrillaux et al. 1973; Stanyon et al. 1992; Stankiewicz et al. 2001).

**Figure 3.**
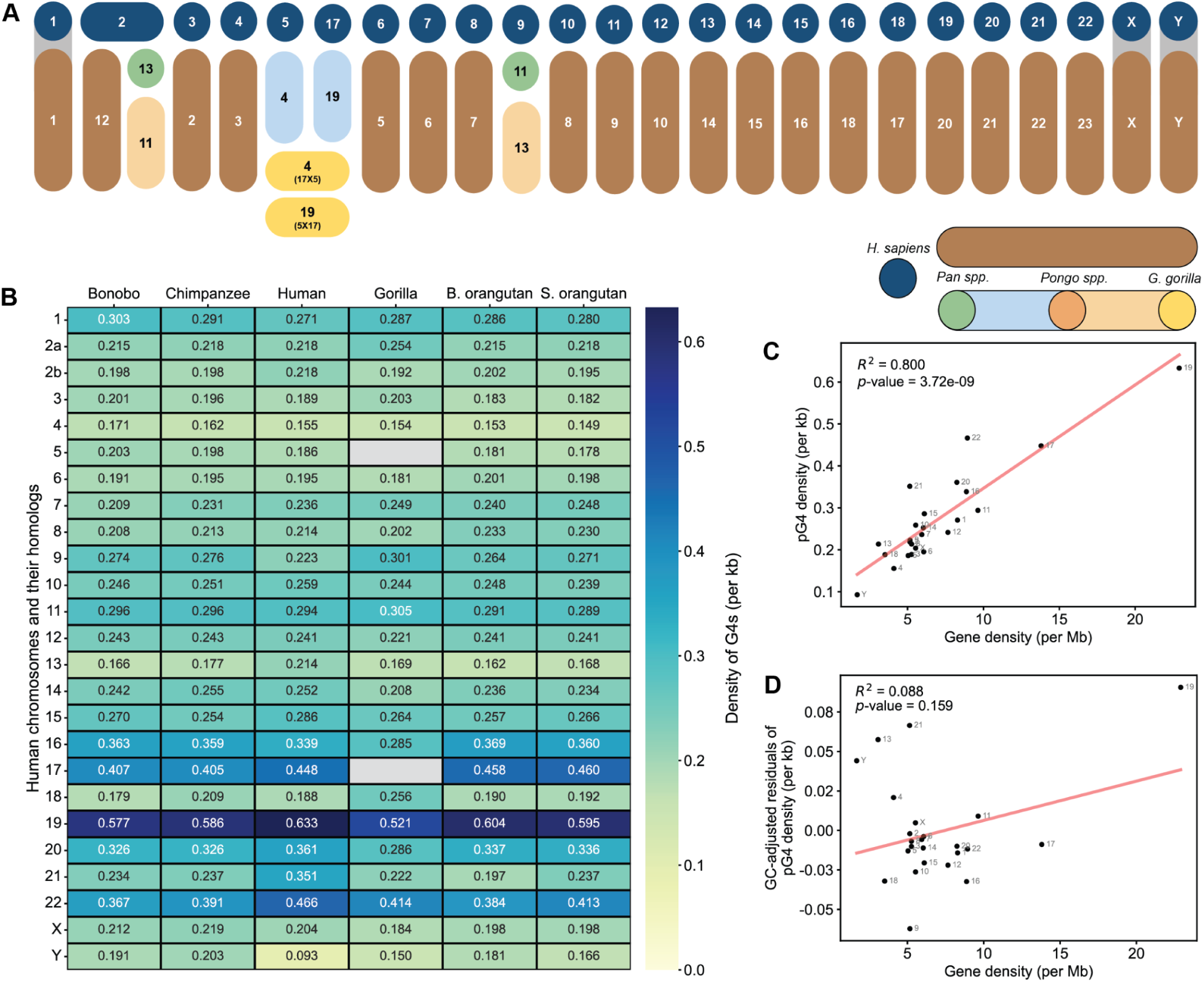
Homologous relationships and pG4 density across great ape chromosomes, and their correlation with gene density in the human genome. (**A**) The map of homologous relationships among great ape chromosomes. Each column represents a homolog, and each row represents a genus or a species. The legend displays the colors used for clades of great apes. Colors that are common to two or more clades indicate shared homology. The numbers over the boxes indicate the chromosome numbers assigned in the T2T assemblies. (**B**) The heatmap of pG4 density with homologous chromosomes (human numbering system is used as a base) as rows and species as columns. Homologs of human chromosomes 5 and 17 for gorilla are excluded from this plot because of the translocation event in gorilla. pG4 density for human chromosome 2 is assigned to two rows—2a and 2b. (**C**) The relationship between pG4 density and gene density across human chromosomes (*p*-value=3.72×10^-9^). (**D**) The same as C, with GC-adjusted residuals for pG4 density and gene density (*p*-value=0.159).

We observed similar pG4 densities for chromosomes homologous between great apes (Fig. 3B). Human chromosomes 17, 19, and 22, as well as their non-human great ape homologs, had high pG4 density (>0.4/kb) compared to that of other chromosomes. We found a positive correlation (*R*^2^ = 0.80) between pG4 density and protein-coding gene density of human chromosomes (Fig. 3C). Genes are known to be GC-rich (Vinogradov 2003; Jaksik and Rzeszowska-Wolny 2012) potentially driving this relationship. While most of the variation in G4 density across chromosomes was explained by their GC content (Fig. S3), gene density still explained a small portion of the variation in G4 density after correcting for GC-content (*R*^2^ = 0.088), even though such association was not statistically significant (Fig. 3D).

### Presence/absence of predicted G4s follows a molecular clock, with many shared and species-specific variants

Using pairwise inter-species genome alignments (see Methods for details), we identified species-specific pG4s *vs.* pG4s shared across different groups of species, forming *evolutionary groups* in our subsequent analyses (Fig. 4A). We found that 271,114 pG4s—approximately one-third of the total number of pG4s in each species—were shared across all species. Closely related congeneric species shared many pG4s as well—197,414 were shared by Bornean and Sumatran orangutans, and 54,741 were shared by bonobo and chimpanzee. Additionally, we found a large number of pG4s shared by Homininae (bonobo, chimpanzee, human, and gorilla sharing 129,730 pG4s) and Hominini (bonobo, chimpanzee, and human sharing 41,585 pG4s). More generally, we found a negative correlation between the number of pG4s shared between any two great ape species and their corresponding divergence times (Fig. 4B).

**Figure 4.**
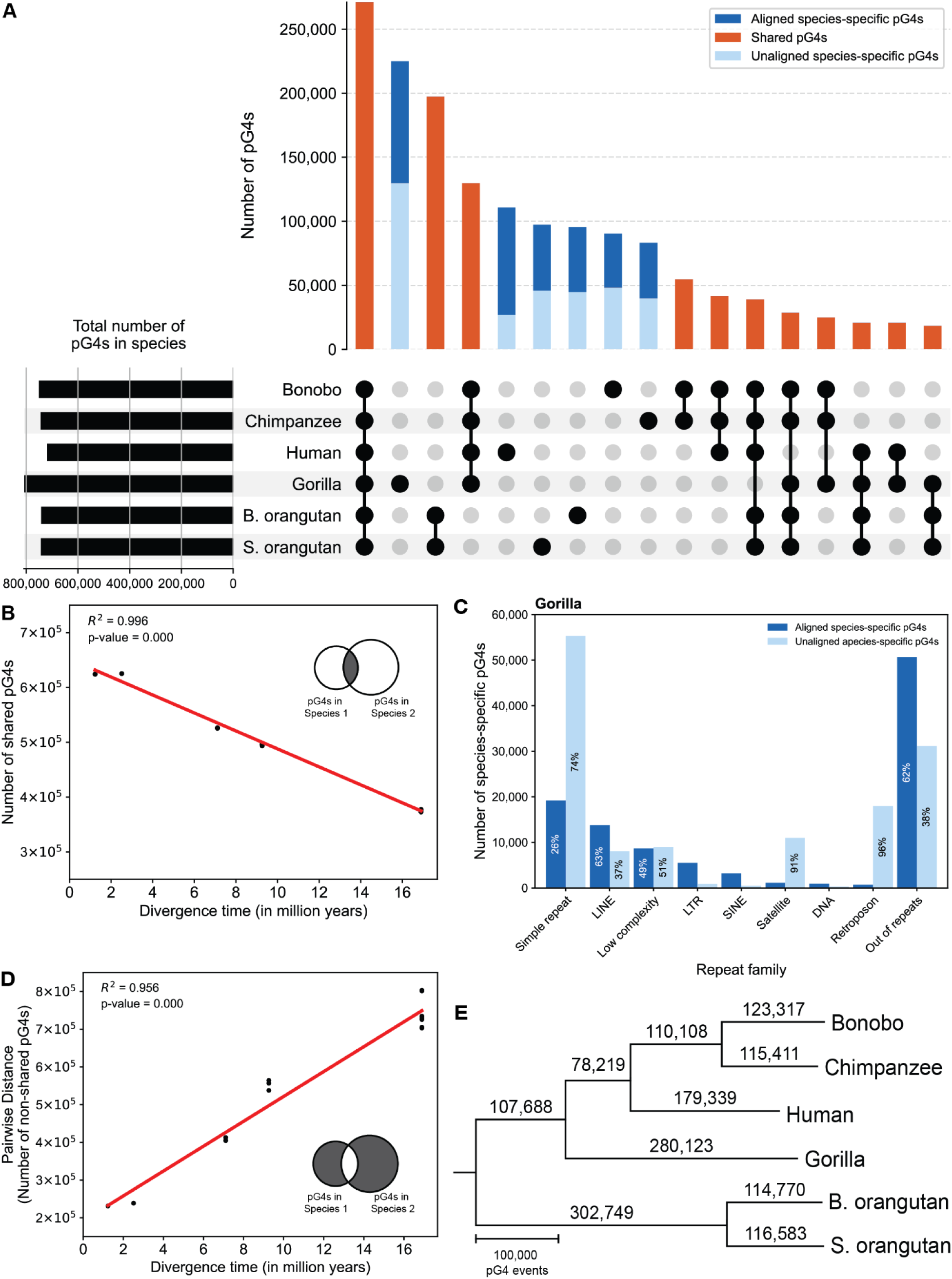
Inter-specific sharing, divergence, and evolutionary dynamics of pG4s in great ape T2T genomes. (**A**) The interspecific sharing of pG4s present at homologous locations of great ape T2T genomes (only the top 17 groups are shown, see Fig. S4 for all the groups). The vertical orange bars represent the number of pG4s shared between different great ape species as shown by the shaded circles, and the vertical blue bars represent the number of species-specific pG4s, with aligned and unaligned pG4s in dark blue and light blue, respectively. The horizontal bars represent the total number of pG4s in each great ape species throughout its genome. (**B**) The correlation between the total number of shared pG4s and divergence time. Black dots represent pairs of species, and the red line represents the best fit. (**C**) The number of aligned vs. unaligned species-specific pG4s in each repeat family—DNA repeats, LINEs, LTRs, low-complexity regions, retroposons, SINEs, satellites, simple repeats, and out-of-repeat regions in gorilla. The percentages inside each bar represent the proportion of aligned/unaligned pG4s in the respective repeat family. (**D**) The correlation between the total number of non-shared pG4s and divergence time. Black dots represent pairs of species, and the red line represents the best fit. The Venn diagram at the bottom right is a schematic explaining pG4s that were considered to calculate the distance between any two species, 1 and 2, based on the number of non-shared pG4s. (**E**) The phylogram inferred as the most parsimonious relationship from pG4 presence/absence data. The branches are scaled to their lengths and represent the number of pG4s on that branch, i.e. the total number of births and deaths of G4 inferred from the parsimony-informative G4s.

Gorilla exhibited the highest number of species-specific pG4s (225,045, or 27% of all gorilla pG4s), followed by human (110,689, or 14%). Chimpanzee had the smallest number of species-specific pG4s (83,173, or 11%). Species-specific pG4s were further divided into *unaligned* and *aligned* (Table 2) because they might arise via different mechanisms. The sequences for unaligned species-specific pG4s were present in one species only and were absent from any pairwise alignments; we hypothesized that such pG4s might have arisen due to lineage-specific repeat and satellite expansions. The sequences for aligned species-specific pG4s were present in pairwise alignments but were annotated as G4s in one species only; we hypothesized that such pGs might have arisen by nucleotide substitutions and/or small insertions and deletions. Across the great apes, 27,006 to 129,860 unaligned species-specific pG4s were identified, constituting 24-58% of their species-specific pG4s. Gorilla had the highest number of unaligned species-specific pG4s (129,860), while the lowest number was observed for human (27,006). Consistent with our hypothesis, the species-specific, and particularly unaligned, pG4s were abundant within the repetitive *vs.* functional regions of the great ape genomes (Fig. S5). Among different repeat classes, simple repeats—frequently present at telomeric and subtelomeric regions (Aksenova and Mirkin 2019)—contained the highest number of unaligned species-specific pG4s, in all species but human (Fig. 4C, Fig. S6). Retroposons, particularly SVA elements, represented the next most frequent class across all species studied, except for orangutans (Fig. S6).

**Table 2.**
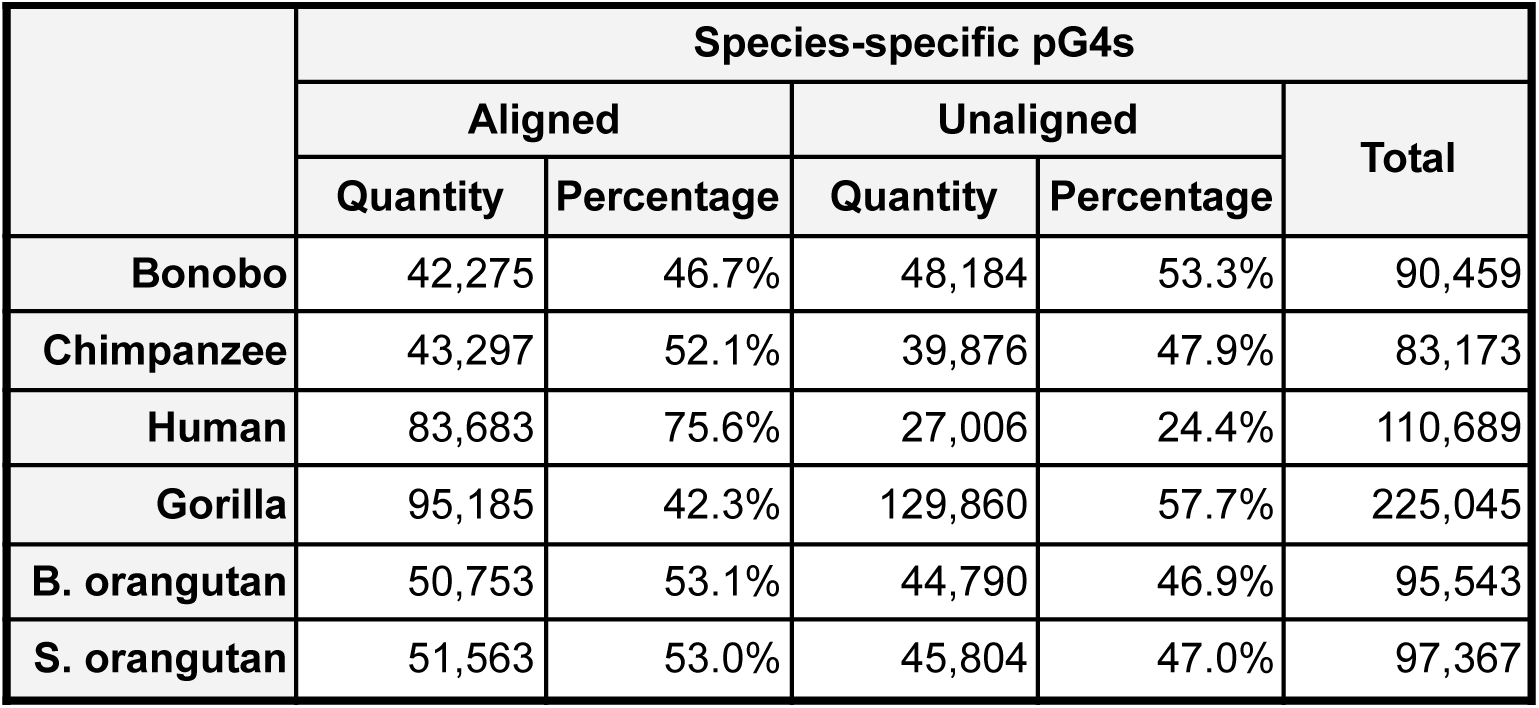
The total number of species-specific pG4s in each great ape species. The numbers and proportions of aligned vs. unaligned species-specific pG4s are also shown. (B. orangutan: Bornean orangutan, S. orangutan: Sumatran orangutan)

|  | Species-specific pG4s |  |  |  |  |
| --- | --- | --- | --- | --- | --- |
|  | Aligned |  | Unaligned |  | Total |
|  | Quantity | Percentage | Quantity | Percentage |  |
| **Bonobo** | 42,275 | 46.7% | 48,184 | 53.3% | 90,459 |
| **Chimpanzee** | 43,297 | 52.1% | 39,876 | 47.9% | 83,173 |
| **Human** | 83,683 | 75.6% | 27,006 | 24.4% | 110,689 |
| **Gorilla** | 95,185 | 42.3% | 129,860 | 57.7% | 225,045 |
| **B. orangutan** | 50,753 | 53.1% | 44,790 | 46.9% | 95,543 |
| **S. orangutan** | 51,563 | 53.0% | 45,804 | 47.0% | 97,367 |

We observed some variation in pG4 distribution across homologous chromosome groups (Fig. S4). The most striking difference was observed in chromosome Y, where pG4 sharing among great apes was minimal, with each species exhibiting a high proportion of species-specific pG4s (39-80%). Additionally, human chromosomes 13 and 21, along with their homologs in non-human great apes, displayed similarly high proportions of species-specific pG4s (21-39% and 22-38%, respectively). Furthermore, in gorilla, chromosome 17 (homologous to human chromosome 18) contained a substantial number of species-specific pG4s, whereas its chromosomal homologs in other species did not (Fig. S4).

We next investigated whether pG4 presence/absence follows the molecular clock. The molecular clock hypothesis postulates that the genetic distance between any two species is proportional to their divergence time (reviewed in (Bromham and Penny 2003)). Here we defined the genetic distance as the number of non-shared pG4s between species, called “the pairwise G4 distance” henceforth. The pairwise G4 distance was calculated by subtracting two times the number of shared pG4s from the total number of pG4s present in the genomes of two species. We found a strong correlation (R² = 0.956) between such distance and divergence time as provided in (Makova et al. 2024), indicating that pG4 presence/absence does indeed follow the molecular clock (Fig. 4D). When we removed the species-specific unaligned G4s from this analysis, we observed an even stronger correlation (R² = 0.997; Fig. S7A). Additionally, we constructed the most parsimonious tree using the presence/absence of pG4s at homologous positions across great ape T2T assemblies (see Methods for details). This tree (Fig. 4E) was consistent with the known species phylogeny (Makova et al. 2024).

### Predicted G4s are enriched and hypomethylated at regulatory regions

We further investigated the enrichment of pG4s for different categories of functional regions in the great ape genomes by computing pG4 densities for each category and comparing them with the respective genome-wide pG4 densities (see Methods). The functional region categories (later called *functional categories*) considered were: promoters, 5’UTR (5’ untranslated regions), protein-coding sequences (CDS), introns, 3’UTRs, enhancers, non-protein coding genes, origins of replication, CpG islands, and repeats (as annotated by RepeatMasker (Smit et al. 2013-2015)). We added to these a category comprising non-functional non-repetitive, presumably neutrally evolving, regions (NFNR). The genic functional categories—UTRs, protein-coding sequences, introns, and non-protein coding genes–were divided into transcribed and non-transcribed strands. Furthermore, pG4s in each category were divided into four evolutionary groups based on their sharing at homologous locations across ape genome assemblies: pG4s shared by great apes, by Hominines, by Hominini, *Pan* or *Pongo* genus-specific pG4s, and species-specific pG4s (Fig. 5, Fig. S8).

**Figure 5.**
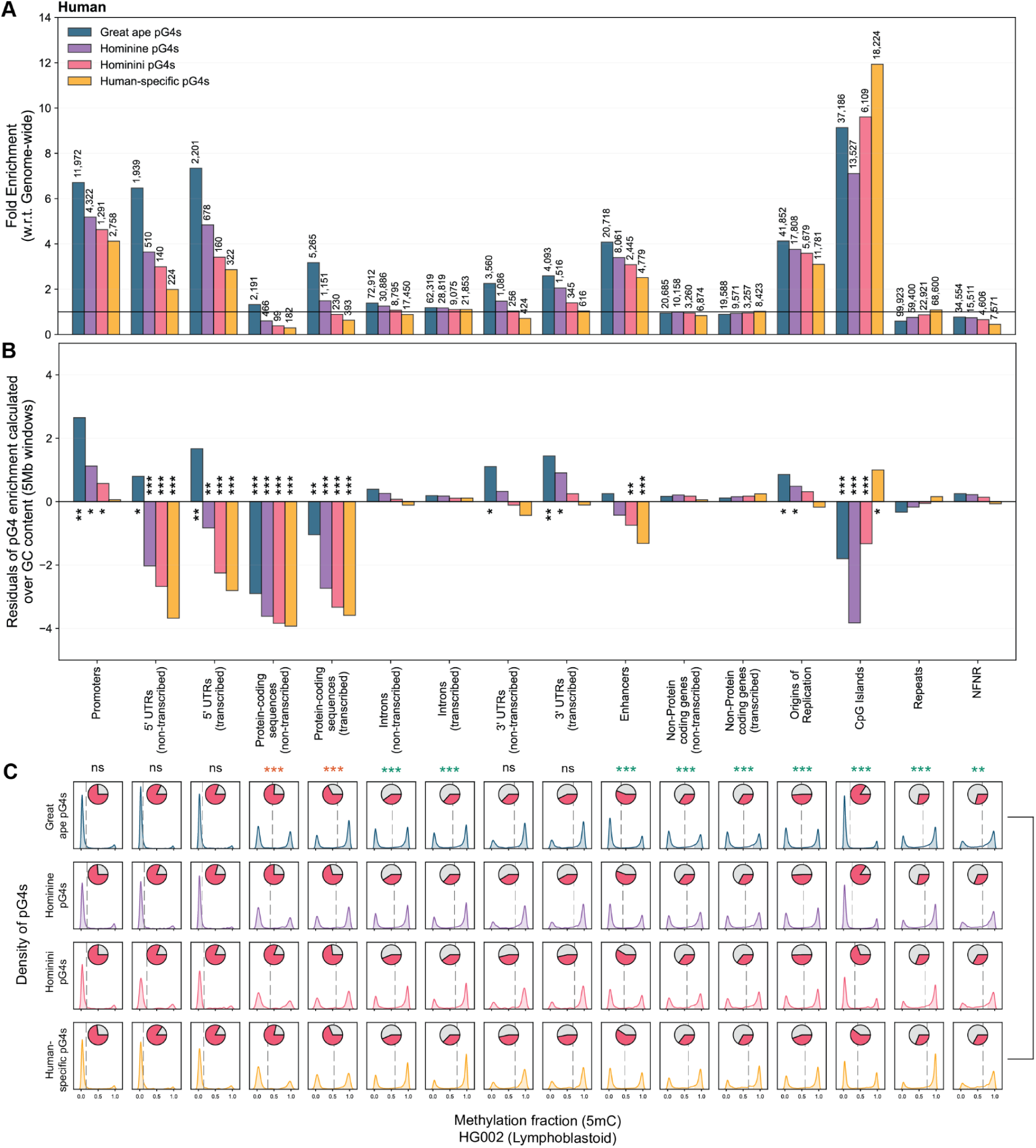
pG4 enrichment and methylation across functional categories and evolutionary groups in the human genome. (**A**) Enrichment levels of pG4s across different categories of functional regions in the human genome. The y-axis displays fold enrichments for each functional category. The bars are color-coded by evolutionary group (great ape, Hominine, Hominini, and human-specific pG4s). The bar heights reflect enrichment values, with numbers atop indicating the total pG4s in each evolutionary group and functional category. (**B**) GC-corrected enrichment (positive residuals indicate enrichment and negative residuals indicate depletion) of pG4s across the same functional categories as in (A). The y-axis displays residuals from the regression of pG4 enrichment on GC content, fitted using 5-Mb non-overlapping genome windows. Stars above/below the bars denote the statistical significance of the GC-corrected enrichment, with significance computed using the percentile rank of residuals within the genome-wide residual distribution of GC content in 5-Mb windows in a two-tailed test (Fig. S9D). (**C**) Methylation distributions (kernel density plots) of pG4s with CpG sites across pG4 evolutionary groups and functional categories for the HG002 cell line. Each pie chart shows the fraction of pG4s with CpG sites (red portion). The rows correspond to pG4 evolutionary groups. The columns match the functional categories in (A) and (B). Significance of methylation differences, calculated using a two-tailed test of proportions, is shown above the columns, highlighting hypomethylation differences between pG4s shared by great apes and human-specific pG4s. Red stars indicate stronger hypomethylation in human-specific pG4s, while green stars indicate stronger hypomethylation in pG4s shared by great apes (ns: not significant, *: *p*-value < 0.05, **: *p*-value < 0.01, ***: *p*-value < 0.001).

In human, pG4s were highly enriched at promoters (4.12 to 6.71 fold), 5’UTRs (1.99 to 7.34 fold), enhancers (2.51 to 4.08 fold), and origins of replication (3.10 to 4.13 fold), with the lowest and highest values listed for the human-specific pG4s and for the pG4s shared across great apes, respectively (Fig. 5A). We also observed pG4 enrichment at CpG islands (7.11 to 11.93 fold). However, here the highest enrichment was observed for the human-specific pG4s and the lowest for Hominine pG4s (Fig. 5A). Protein-coding sequences displayed both strand– and group-specific enrichment patterns. Specifically, pG4s were highly enriched on the transcribed strand for both great ape and Hominine shared pG4s (1.49 and 3.18-fold, respectively) and moderately enriched on the non-transcribed strand for great ape shared pG4s (1.32-fold) (Fig. 5A). In contrast, 3’UTRs exhibited enrichment on both strands, with the greatest enrichment observed in great ape shared and Hominine pG4 evolutionary groups (1.47 to 2.59-fold; Fig. 5A). pG4s at non-protein coding genes, introns, repeats and NFNR were either at the genome-level enrichment or depleted (Fig. 5A). Non-human great apes followed similar trends of pG4 enrichment across functional categories (Fig. S8A,D,G,J,M).

Given the correlation between G4 density and GC content (Fig. S9B), we corrected for the latter regressing pG4 fold enrichment over GC content genome-wide and considering residuals, which capture enrichment not explained by GC content (Fig. 5B, Fig. S8B,F,H,K,N; see Methods for details). After such correction, pG4s shared across great apes were still significantly enriched at promoters, 5’ and 3’ UTRs (both strands), and origins of replication (only for human). Other evolutionary groups of pG4s in some instances switched from enrichment to depletion after the GC correction. All pG4s evolutionary groups were significantly depleted at protein-coding sequences, with particularly strong depletion for the non-transcribed strand. Additionally, after the GC correction, CpG islands were significantly depleted in pG4s, except for species-specific pG4s, which still showed enrichment.

Using the genome-wide methylation data obtained from long-read sequencing of the human genome (Gershman et al. 2022), we evaluated the methylation status of pG4s in a lymphoblastoid cell line (HG002 cell line) and a hydatidiform mole cell line (CHM13). We performed a similar analysis with the cell lines from which the non-human great ape T2T assemblies originated—fibroblast cell lines for bonobo, gorilla, Sumatran orangutan, and Bornean orangutan, and a lymphoblastoid cell line for chimpanzee (Yoo et al. 2024). This allowed us to predict G4 formation *in vivo*, as methylated sequences were shown to be less likely to form G4 structures (Halder et al. 2010; Mao et al. 2018). We examined the 5-methylcytosine methylation profiles of pG4s that contain CpG sites within the functional categories in the cell lines analyzed (Fig. 5C, Fig. S11A). Following a previous study (Gershman et al. 2022), we defined hypomethylated and hypermethylated pG4s as having methylation fraction lower than 0.2 and higher than 0.8, respectively. In some functional categories, including introns, 3’UTRs, non-protein coding genes, repeats, and NFNRs, less than half of pG4s had CpG sites (Fig. 5C, Fig. S8C,G,I,L,O). This suggests that G4s in these regions are regulated in ways other than 5mC methylation.

We made three important observations about hypomethylated pG4s, which have a high probability to form *in vivo*. First, in HG002, G4 methylation levels were low across all pG4 evolutionary groups at 5’UTRs (both transcribed and non-transcribed strands), promoters, enhancers, and CpG islands—suggesting G4 formation in these functional categories. This was true for other great ape cell lines as well (Fig. S8C,G,I,L,O). Second, across all great apes, pG4s shared by great apes consistently exhibited a significantly higher proportion of hypomethylation (*p*-value < 0.001, two-tailed test of proportions) compared to species-specific pG4s in most functional categories—except for promoters, UTRs, and protein-coding sequences (significance stars in Fig. 5C and Fig. S10). In promoters and 5’ UTRs, this difference was significant in chimpanzee, bonobo, and gorilla, while in 3’ UTRs, it was significant in gorilla alone. Third, whereas the CHM13 cell line exhibited a methylation profile similar to that for HG002, the hypomethylated fraction of pG4s shared among great apes in CHM13 was significantly higher than that in HG002 in all functional categories except for promoters, CpG islands, protein-coding sequences and 5’ UTRs (Fig. S11C). Broadly, this observation is consistent with lower levels of methylation in tumor compared to normal cells (Hoffmann and Schulz 2011; Besselink et al. 2023) and suggests a stronger activation of G4s in the CHM13 cell line as compared to the HG002 cell line.

## DISCUSSION

Using recently released great apes T2T genomes, we conducted a detailed analysis of occurrence of predicted G4s—focusing on their evolution and enrichment in different functional regions. To predict G4s based on genome sequence information, we combined two commonly used G4 prediction algorithms: pqsfinder (Hon et al. 2017) and G4Hunter (Bedrat et al. 2016). Our pipeline identified many new pG4s, which were previously unnoted likely due to incomplete sequence information, and created a comprehensive pG4 catalog for human as well as non-human great apes, labeling predictions as shared among different evolutionary clades or species-specific. With its focus on great apes, our study complements previous studies of G4 evolution, which considered larger evolutionary distances, e.g., spanning different kingdoms of life (Vannutelli et al. 2023) or early diverging eukaryotic clades (Wu et al. 2021). Previously G4 occurrence was studied only for three human T2T chromosomes (Bohálová et al. 2021; Brázda et al. 2022; Dobrovolná et al. 2024) and was not analyzed for great ape chromosomes. Using pairwise alignments, G4 predictions, and homology information of great ape chromosomes, we were able to predict species-specific G4s and G4s shared across different species clades using a graph-based approach. Chromosomal-level analysis showed that the density of G4s is linearly correlated with the gene density, and the high GC content of the gene-enriched chromosomes explains most of this pattern. Moreover, using methylation data as a proxy for G4 formation in the cell, we were able to predict G4 formation in different types of functional regions, such as promoters and origins of replication.

### Newly discovered pG4s in great apes

The number of pG4s generated by our pipeline for the human genome was higher (by 32,499-52,878) than previous estimates, which reported a total number of G4s between 716,310 and 736,689 (Chambers et al. 2015; Marsico et al. 2019; Tu et al. 2021). This increase is explained by our use of the T2T genome assembly and by our prediction of not only standard but also bulged G4s. We found that the newly resolved regions in the T2T genomes of all great apes were enriched for pG4s, consistent with overall enrichment for non-B DNA in such regions (Smeds et al. 2024). In human, we detected a particular enrichment of G4s at the acrocentric chromosomes. In non-human great apes, we detected newly resolved G4-dense regions mostly in the metacentric chromosomes (Fig. S2).

### Evolution of pG4 occurrence in great apes

At the genome level, we found that pG4s evolve following the molecular clock hypothesis (Fig. 4D), indicating a consistent rate of G4 divergence over time. This suggests that, globally, pG4s evolve with similar rates and under similar selective constraints in the great ape lineages. Previous studies suggested that G4s are subject to purifying selection in several functional categories of the human genome, with different levels of constraint depending on the regions considered (Guiblet, DeGiorgio, et al. 2021). Our study suggests that such constraints might be similar across the studied ape species, although a more detailed investigation is warranted. The strong negative correlation we found between the number of pG4s shared between species and the species divergence time (Fig. 4B) is also supportive of their evolution following the molecular clock hypothesis.

Our study also highlighted species-specific pG4s. In particular, we discovered different evolutionary and genomic distribution patterns between aligned and unaligned species-specific pG4s. The aligned species-specific pG4s followed a rather strict molecular clock (Fig. S7B), consistent with them originating via nucleotide substitutions and/or small insertions/deletions, which are known to follow the molecular clock. In contrast, the unaligned species-specific pG4s followed a more relaxed molecular clock (Fig. S7C), and were located predominantly in the repetitive regions of the great ape genomes (which were deciphered in the T2T assemblies), in agreement with a recent study (Smeds et al. 2024). We found that pG4s are common at simple repeats (including telomeres), satellites, and retroposons (Fig. 4C, Fig. S6), which were previously suggested to expand rapidly and frequently in a genus– and/or species-specific manner (Cechova et al. 2019; Hoyt et al. 2022; Makova et al. 2024; Yoo et al. 2024).

For the first time, we also showed that for pG4s in different functional categories, there is a significant difference between the proportion of hypomethylated G4s, depending on their level of sharing across the great apes. In particular, in most functional region categories, shared G4s had higher levels of hypomethylation, and thus are more likely to form, as compared to species-specific G4s. The opposite was true for human protein-coding genes, where shared pG4s were more methylated than species-specific ones. Additional studies are needed to investigate the potential causes of these relationships.

### Functional regions display a bias towards G4s

Our study corroborates a correlation between G4 density and GC-content that was observed previously (Wu et al. 2021). Moreover, our functional enrichment study, in line with previous studies (Vannutelli et al. 2022), provides evidence that the prevalence of G4s cannot be solely attributed to the GC content of any functional category. Some functional categories, such as promoters and origins of replication, show a significant G4 enrichment even after GC correction. However, some other functional categories are no longer significantly enriched for G4s after a GC correction.

#### Promoters

The enrichment of G4s we found at promoters corroborates previous studies (Huppert and Balasubramanian 2007; Lago et al. 2021; Wu et al. 2021). We also found that G4s located at promoters were hypomethylated, indicating significant G4 activation at these regions of the genome. This result is consistent with results by Wu and colleagues (2021), who observed significantly lower methylation levels at G4s located within 2 kb upstream of genes *vs*. the rest of the genic regions in mammals and insects. These observations are in agreement with the critical role G4s play in the promoter regions during gene regulation (e.g., (Huppert and Balasubramanian 2007; Tian et al. 2018)).

#### CpG islands

GC-rich CpG islands were highly enriched in G4s—particularly before GC correction and at human-specific CpG islands even after GC correction. CpG islands are often found at promoters, i.e., almost one-fourth of all promoter regions in our study overlapped with CpG islands. Studies have shown regulatory roles of G4s in CpG islands (Bay et al. 2017). Since CpG islands are typically hypomethylated, they provide a favorable environment for activated G4s, particularly at promoter regions (Mao et al. 2018).

In gorilla, we found a notable enrichment of species-specific G4s at CpG islands after correcting for GC content (Fig. S8H). Interestingly, unlike human-specific pG4s at CpG islands, these G4s were hypermethylated (Fig. S8I). A similarly strong pattern of hypermethylation appeared in gorilla repeat regions, suggesting that gorilla may have experienced repeat expansions (Makova et al. 2024; Yoo et al. 2024) leading to the formation of new G4 structures (Fig. S8I). The difference might also be due to different cell lines analyzed for human (lymphoblastoid) vs. gorilla (fibroblast). The increased methylation at these sites could represent an evolutionary response to suppress potential G4 activation, reflecting a regulatory adaptation in gorilla.

#### Enhancers

Enhancers also showed both enrichment and hypomethylation of G4s, though generally to a lesser degree than what was observed in promoters, across all great apes. This aligns with previous studies (e.g., such as (Hegyi 2015)), which have proposed that G4s play a role in promoter-enhancer interactions, with G4s forming partly in the promoter and partly in the enhancer. Another study (Williams et al. 2020) further supported the possibility of these G4-mediated interactions, suggesting that enhancers with long G4-forming sequences could be instrumental in such regulatory contacts. However, after correcting for GC content, the observed G4 enrichment in enhancers was often reduced, suggesting that GC content largely accounts for the presence of G4s in these regions. Therefore, while our observations do not directly confirm these models, they suggest that G4s may act as secondary players in enhancer regulation, contingent upon GC-rich contexts.

#### UTRs

Prior to GC correction, pG4 enrichment was observed in both 5’ UTRs and 3’ UTRs across all great apes, although the enrichment was less pronounced in 3’ UTRs. This observation is in support of previous studies (Huppert et al. 2008). However, similar to other functional groups, the enrichment decreased with the level of sharing—going from the most shared evolutionary group of great apes to the human-specific ones. After GC correction, however, enrichment patterns varied by evolutionary group. For both 5’UTRs and 3’UTRs, pG4s shared across all great apes showed significant enrichment, while other evolutionary groups in 5’UTRs showed a depletion of pG4s. This consistent pattern across great apes suggests that while pG4 structures may be functionally relevant within UTRs, pG4s that are highly conserved across species are favored. Further investigation is needed to understand the selective mechanisms underlying this conservation at pG4s shared by great apes. Whereas G4s at 5’UTRs were mostly hypomethylated, G4s at 3’UTRs showed a dominant hypermethylated fraction. Additionally, nearly three-quarters of G4s in 5’UTRs contained CpG sites, whereas fewer than half of G4s in 3’UTRs did. This disparity, evident across all great apes, suggests that G4s play different roles in 5’ *vs*. 3’ UTR biology. Whereas 5’UTR G4s might influence transcriptional regulation (e.g., they may act as transcriptional barriers, potentially stalling transcription and representing considerable roadblocks for transcriptional machinery), 3’UTR G4s could contribute to post-transcriptional processes. Thus, our study suggests a more nuanced functional landscape for G4s at UTRs, significantly augmenting previous studies (Bugaut and Balasubramanian 2012; Qi et al. 2021).

#### Protein-coding sequences

In line with a previous study (Guiblet, DeGiorgio, et al. 2021), we found that pG4 enrichment, albeit modest, had a strand-specific pattern, in the protein-coding sequences—with the transcribed strand having a higher enrichment as compared to the non-transcribed strand, for shared pG4s. This is in line with the hypothesis that G4s are not favored at the mRNA level (Huppert and Balasubramanian 2005). We found that protein-coding sequences, although known to be GC-rich, displayed a significant G4 depletion after correcting for GC content (Huppert and Balasubramanian 2005; Varizhuk et al. 2017), across all evolutionary groups, and at both transcribed and non-transcribed strands. Thus, G4 formation is disfavored at these regions. Interestingly, across all great apes, species-specific G4s in the non-transcribed strand of the protein-coding sequences had significantly higher hypomethylation as compared to G4s shared by great apes. Therefore, it can be hypothesized that evolutionarily new G4s have a tendency to be active in the non-transcribed strand, which might result in G4 formation in mRNA. Alternatively, such G4s might not have had enough time to become silenced by methylation. This interesting observation should be investigated further.

#### Replication origins

Our analysis revealed an enrichment of G4s at human replication origins. While this enrichment decreased after correcting for GC content, there remains a positive trend that suggests a connection between G4s and replication origins, particularly in more highly conserved G4s as opposed to human-specific ones. This trend hints at a functional relationship between G4 structures and replication initiation suggested in previous studies (Prioleau 2017; Prorok et al. 2019). Due to the rapid evolution of origins of replication (Massip et al. 2019), we were unable to study their enrichment and methylation status in other great apes.

### Study limitations, future directions, and conclusions

This study, based exclusively on sequence data, underscores the necessity for further experimental validation to confirm G4 formation *in vivo* in great apes and to enhance our understanding of their evolutionary dynamics. Several factors were not considered in our analysis, which may influence G4 prediction accuracy. These include the loop-base composition (Puig Lombardi et al. 2019), the presence of uneven G4 motifs (Maity et al. 2020), and the impact of adenine repeats on G4 stability (Chen et al. 2017). Incorporating these factors into future studies could refine our G4 predictions.

Future research could also revisit some of the limitations of our study. First, our G4 repertoire was defined as a non-overlapping set, selecting the most stable G4s (based on the highest pqsfinder scores) in each genomic interval. However, under physiological conditions, multiple G4 conformations can form, and sequence data alone cannot fully capture this complexity. Experimental approaches (e.g., permanganate/S1 footprinting with direct adapter ligation and sequencing (Lahnsteiner et al. 2024)) will be essential to identify these dynamic G4 structures *in vivo*. Second, for the sharing profile of G4s, we only considered those that fully overlapped with alignment blocks, which may have led to an underestimation or overestimation of shared or species-specific G4s, respectively, among great apes. While these excluded G4s likely represent a small portion of the overall repertoire, they could form a distinct group deserving of separate analysis. Third, we used a quadratic regression model to correct for GC content when assessing G4 enrichment. While this worked satisfactorily for our data, a more sophisticated model (e.g., logistic growth model) may better reflect the biological relationship between G4 formation and GC content and provide a more accurate understanding of how G4 enrichment varies with GC content. Finally, our analysis did not consider the predicted stability of G4s. Incorporating such information in future analyses could be informative, as, for instance, a previous study provided evidence of depletion of thermodynamically stable G4s in the genomes of multiple species (Puig Lombardi et al. 2019). Taken together, our whole-genome catalogue and analyses of G4s in humans and other great apes open new avenues for comparative research of G4 evolution.

## MATERIALS AND METHODS

### Predicting G4s using pqsfinder and G4Hunter

Genome FASTA files for six great ape species (version 2.0)—*Homo sapiens* (NCBI RefSeq: GCF_009914755.1), *Pan troglodytes* (NCBI RefSeq: GCF_028858775.2), *Pan paniscus* (NCBI RefSeq: GCF_029289425.2), *Gorilla gorilla* (NCBI RefSeq: GCF_029281585.2), *Pongo abelii* (NCBI RefSeq: GCF_028885655.2), and *Pongo pygmaeus* (NCBI RefSeq: GCF_028885625.2)—were downloaded from NCBI. Each of these FASTA files was divided by chromosome, and only the primary haplotype was used. pqsfinder (dockerized at https://github.com/kxk302/PqsFinder_Docker/tree/main) was employed on all files with the following settings: `overlapping=True`, `maxLength=50` and `minScore=30`. The pG4s that satisfied the regular expression motifs of standard and bulged G4s, contained more than two tetrads, had a pqsfinder score of 40 or higher, and had an absolute G4Hunter score (adapted from https://github.com/AnimaTardeb/G4Hunter/blob/master/G4Hunter.py) greater than 1.5 were retained. Following this, the pG4s in the same region (i.e. between a certain start and end position) were grouped together and only the pG4s with the highest pqsfinder score were selected for further analyses. The final dataset consisted of a non-overlapping set of pG4s on both strands for all the chromosomes of the great apes.

### pG4 density and distribution across great ape genomes

The pG4 density for the great ape chromosomes was calculated by dividing the number of pG4s by the length of the respective chromosome in each species. Inter-species chromosomal homolog mapping was inferred from the Comparative Genome Viewer from NCBI, which utilizes pairwise alignments between the genomes of the great apes. To study the distribution of pG4s, chromosomes were divided into 100-kb windows, and the number of pG4s in each window was tabulated. The density for each window was then calculated by normalizing the number of pG4s in that window against the window with the highest number of pG4s on the same chromosome.

### Pairwise genome alignments using LASTZ between great ape homologous chromosomes

The FASTA files used for the prediction of G4s were also utilized for pairwise alignments of homologous chromosomes among the great apes (see previous section), as illustrated in Fig. 3A. To account for the fusion event in human, all the great ape chromosomes homologous to human chromosome 2a and 2b, were aligned to the whole human chromosome. In addition, pairwise alignments were performed among the non-human great ape chromosomes corresponding to 2a and 2b. Similarly, to account for the translocation event in the gorilla involving the human homologs of chromosomes 5 and 17, the pairwise alignments for these homologs included both gorilla chromosomes 4 and 19. For all human homologs other than 2, 5, and 17, a total of 15 pairwise alignments were performed for each homolog across the six great ape species. Alignments were performed using LASTZ (Harris 2007) with the settings `--notransition` and `--allocate:traceback=1.5G`. The human-chimp.v2 scoring matrix (https://genomewiki.ucsc.edu/index.php/Hg19_conservation_lastz_parameters) was used for these pairwise alignments.

### Connected graphs for shared and species-specific G4s

Predicted G4s were mapped onto the pairwise alignments and classified as shared or species-specific using our custom-designed pipeline—MAP-SEA (Mapping And Prediction of Shared Elements in Alignments). The mapping was performed using the BEDTools suite (Quinlan and Hall 2010; Dale et al. 2011). A pG4 was classified as “shared” if its start and end positions were within 3 base pairs (bp) of each other in the pairwise alignments, allowing for up to 3 bp extensions or truncations at either or both ends. Shared G4s from all pairwise alignments across all chromosomes of the great apes were identified as connecting edges, which were subsequently grouped to form connected graphs of shared pG4s across all species. Creating connected graphs removed the problem of double counting a particular pG4, which can arise from repetitions in alignments. pG4s in alignments that showed no sharing and those not included in the alignments were categorized as species-specific pG4s. Each shared group of pG4s, along with species-specific pG4s, was assigned a unique identifier. The connected graphs were collectively stored as a single dataframe encompassing all pG4s. A diagram of this approach is shown in Fig. S12.

A presence/absence matrix was calculated from the dataframe above for downstream analyses. To identify whether a pG4 is present or absent in a species, duplicated pG4s in any single species were counted only once. This caused a reduction in total pG4 counts, as shown by the horizontal bars in Fig. 4A. To visualize the sharing profile, the dataframe was represented as an upset plot (Lex et al.) using the Python package UpSetPlot (https://upsetplot.readthedocs.io/en/stable/index.html).

### Reconstructing the phylogenetic relationship with maximum parsimony

The pG4s presence/absence data at orthologous locations was used for inferring the most parsimonious phylogram for the great apes. The data was converted into a NEXUS format file following the guidelines provided in the PAUP (Swofford 2003) (https://paup.phylosolutions.com/) quick start tutorial. A maximum parsimony heuristic search was conducted designating the orangutans as the monophyletic outgroup. The resulting best-rooted tree was saved in Newick format and subsequently visualized using MEGA (Stecher et al. 2020).

### Functional annotations for humans and other great apes

Genome GFF files for six great apes were downloaded from NCBI. Protein-coding genes with the longest corresponding mRNA, CDS, and exon entries were extracted (using the snippet at https://gist.github.com/karolpal-jr/48213a0a65475e44f708d5d815127bc3). Introns were inferred from the mRNA and exon data, representing the non-exonic regions of mRNA. The 5’ and 3’ UTRs were determined using CDS and exon entries, representing the non-CDS regions of exons at the 5’ and 3’ ends, respectively. Promoters were defined as 1 kb upstream of the gene start sites. All the genes, excluding pseudogenes, not encoding an mRNA, were identified as non-protein coding genes. We used repeat annotations generated with RepeatMasker (Smit et al. 2013-2015).

For the human genome, core and stochastic origins of replication annotations in the hg38 genome were downloaded from (Akerman et al. 2020), and the UCSC Genome Browser’s liftOver tool was used to convert the coordinates to the CHM13 assembly. Unmasked CpG island annotations were downloaded from the UCSC table browser for the CHM13v2.0 genome. Human 5mC methylation data for CHM13, generated using ONT data, was downloaded in BED format (Gershman et al. 2022; Rhie et al. 2023). All annotations were converted to BED format for further analysis.

CpG annotations for the non-human great apes were obtained with the same algorithm used to calculate CpG island annotations for humans (Gardiner-Garden and Frommer 1987; Miklem and Hillier 2022). Enhancer annotations for the non-human great apes were obtained with the UCSC liftOver tool, using chain files derived from wfmash (Guarracino et al. 2021) all-to-all alignments of primates and human haplotypes (CHM13, HG002 mat/pat, GRCh38) (Yoo et al. 2024).

Methylation data for human HG002—mapped to CHM13 using liftOver tool—and the non-human great apes were adapted from (Yoo et al. 2024). NFNR regions in the human genome were defined as those that do not overlap with genes, enhancers, promoters, origins of replication, and repeats. A similar definition was applied to the non-human great apes; however, due to the lack of annotations for origins of replication in their genomes, and because of the rapid evolution of origins of replication among species (Massip et al. 2019), NFNR regions were calculated without excluding overlaps with origins of replication.

### G4 enrichment and average methylation profiles for functional regions across great ape genomes

Using the dataframe generated from the connected graphs, pG4s were grouped based on their sharing across great apes. In each evolutionary group, pG4s entirely located within `(f=1.0)` each functional category were identified using *bedtools intersect*. In addition, to account for the strand-specificity of pG4s in genic elements—introns, protein-coding sequences, and UTRs—pG4s in these functional categories were divided into transcribed and non-transcribed, using the `-S` and `-s` flags in *bedtools intersect*, respectively. Subsequently, the fold enrichment of pG4s for each functional category within an evolutionary group was calculated by dividing the fraction of pG4s in that category for the group by the proportion of the category’s total length relative to the entire genome length. In symbols, for a given functional category *X* and evolutionary group *Y*, the fold enrichment (*FE*) was calculated as:

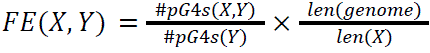

To calculate GC-corrected G4 enrichment, the human genome was divided into 5-Mb windows using bedtools makewindows. GC content for each window was determined with *bedtools nuc*. A second-order polynomial regression was then used to model the relationship between G4 fold enrichment and GC content. We tested a range of window sizes—10 kb, 50 kb, 100 kb, 500 kb, 1 Mb, 5 Mb, 10 Mb, 50 Mb, and 100 Mb—and found that the resulting regression coefficients’ estimates were consistent across window sizes (Fig. S9C). Using the regression model of G4-enrichment against GC content fitted using 5-Mb windows, we calculated the residuals for each functional category within an evolutionary group based on its GC content and observed G4 fold enrichment. The same procedure was applied to calculate the GC-corrected G4 enrichment for non-human great apes. The significance of enrichment, post-GC correction, was calculated using the percentile rank of residuals within the genome-wide residual distribution in a two-tailed test.

The average methylation fraction for each CpG-containing pG4 was calculated by taking the mean of the methylation fraction—the fraction of samples methylated at a given CpG site—across all CpG sites present in the pG4. A kernel density estimate plot was used to visualize the distribution of the average methylation fraction for CpG-containing pG4s in different functional categories across the genomes of great apes. For inter-group differences across all functional categories, the significance of the difference in the proportions of hypomethylation (methylation fraction <0.2) between pG4 evolutionary groups was computed using a two-tailed test of proportions. To account for multiple testing relative to each functional category, *p*-values were Bonferroni-corrected.

## DATA & CODE AVAILABILITY

All data and in-house scripts generated for this paper are available on github: https://github.com/makovalab-psu/GreatApeT2T-G4s.

## AUTHOR CONTRIBUTIONS

S.K.M.: Conceptualization, Data curation, Formal analysis, Investigation, Methodology, Software, Visualization, Writing–original draft. F.C.: Validation, Writing–review & editing. K.D.M.: Conceptualization, Funding acquisition, Methodology, Project administration, Resources, Supervision, Validation, Writing–review & editing.

## ACKNOWLEDGMENTS

We are grateful to Xinru Zhang, Linnéa Smeds, Edmundo Torres-González, Jacob Sieg, and Christian Huber for discussions of the results and useful suggestions. Kaivan Kamali dockerized the pqsfinder package for G4 prediction using fasta files. Karol Pál provided code for extracting protein-coding genes from the great ape gene annotation files. Yong Hwee Eddie Loh, Soojin Yi, and Ryan Son provided useful insights for analyzing methylation data.

