## Supplementary Materials for "Evolutionary Dynamics of G-Quadruplexes in Human and Other Great Ape Telomere-to-Telomere Genomes"

#### SUPPLEMENTARY TABLES

**Table S1. The percentage of pG4s retained at each step of the G4 discovery pipeline.**

<table>
<tr><th colspan="3" rowspan="2">G4 discovery pipeline steps</th><th colspan="2">Human T2T</th></tr>
<tr><th>Number of predicted G4s</th><th>Proportion of predicted G4s</th></tr>
<tr><td colspan="3">G4 prediction using pqsfinder<br>(score ≥ 30 & overlapping=True)</td><td>137,185,107</td><td>100%</td></tr>
<tr><td rowspan="3">Filtering criteria</td><td>1</td><td>Satisfy biological G4 motifs</td><td>14,207,473</td><td>10.4%</td></tr>
<tr><td>2</td><td>pqsfinder score ≥40,<br>G4Hunter score ≥1.5</td><td>10,327,512</td><td>7.5%</td></tr>
<tr><td>3</td><td>Non-overlapping G4s</td><td>769,188</td><td>0.6%</td></tr>
</table>

**Table S2. The top 20 overlaps between different functional categories in the human genome with respect to the whole genome.**
NPC: Non-protein coding sequences, ORs: origins of replication, CpGi: CpG islands.

| Number of Bases | Genome % | Intersection Type |
|---:|---:|---|
| 487,128,494 | 15.63% | Intron, Repeats |
| 216,643,636 | 6.95% | NPC, Repeats |
| 32,363,175 | 1.04% | Intron, NPC |
| 29,992,000 | 0.96% | Intron, NPC, Repeats |
| 23,418,590 | 0.75% | ORs, Repeats |
| 20,656,651 | 0.66% | Intron, ORs |
| 16,152,395 | 0.52% | Intron, ORs, Repeats |
| 11,799,583 | 0.38% | Intron, Enhancer |
| 8,648,493 | 0.28% | CpGi, Repeats |
| 7,680,942 | 0.25% | Enhancer, Repeats |
| 6,019,191 | 0.19% | 3'UTR, Repeats |
| 5,925,947 | 0.19% | Intron, Enhancer, Repeats |
| 5,885,248 | 0.19% | NPC, ORs, Repeats |
| 3,980,429 | 0.13% | NPC, ORs |
| 3,374,245 | 0.11% | Intron, CpGi |
| 3,145,078 | 0.10% | Promoter, Repeats |
| 2,744,682 | 0.09% | Enhancer, NPC |
| 2,583,830 | 0.08% | NPC, CpGi, Repeats |
| 2,413,059 | 0.08% | Intron, CpGi, Repeats |
| 2,079,463 | 0.07% | Enhancer, NPC, Repeats |

### SUPPLEMENTARY FIGURES

**Figure S1. pqsfinder and G4Hunter score distributions from a thresholding run.** (A) pqsfinder and (B) G4Hunter score distributions for predicted G4s in the human T2T genome with relaxed parameters: i.e. pqsfinder score ≥30 and G4Hunter score ≥0. The vertical red line represents the chosen threshold for the scoring algorithm.

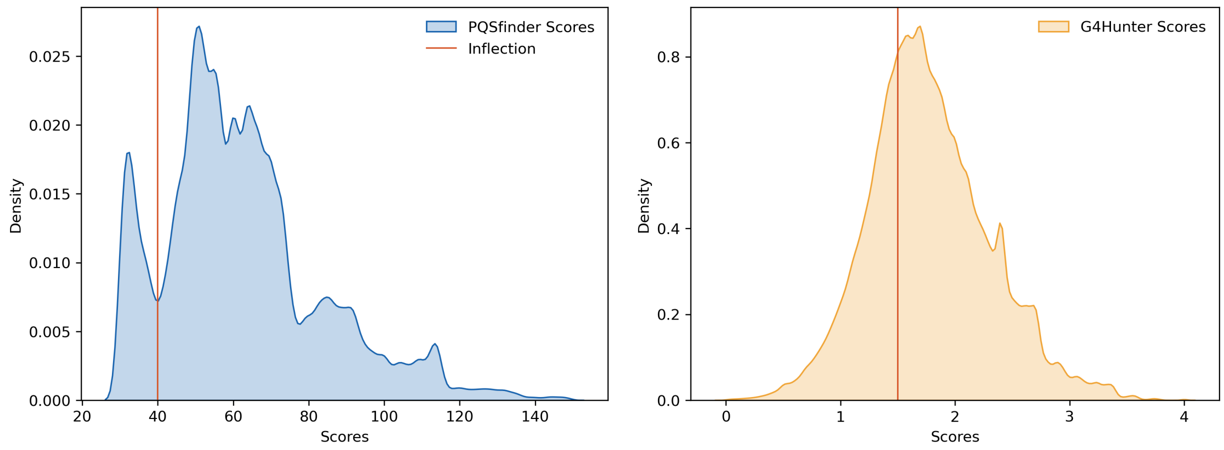

**Figure S2. G4s predicted in the newly resolved regions of non-human great ape T2T genome assemblies.** The number of G4s predicted in the newly resolved regions (per million bases) of the (A) bonobo, (B) chimpanzee, (C) gorilla, and (D) Sumatran orangutan T2T genomes, as compared to the previous assemblies, across all chromosomes. The horizontal red line indicates the number of pG4s per million bases across the human T2T genome. The numbers on the top of each bar show the number of pG4s in the newly resolved regions on each chromosome.

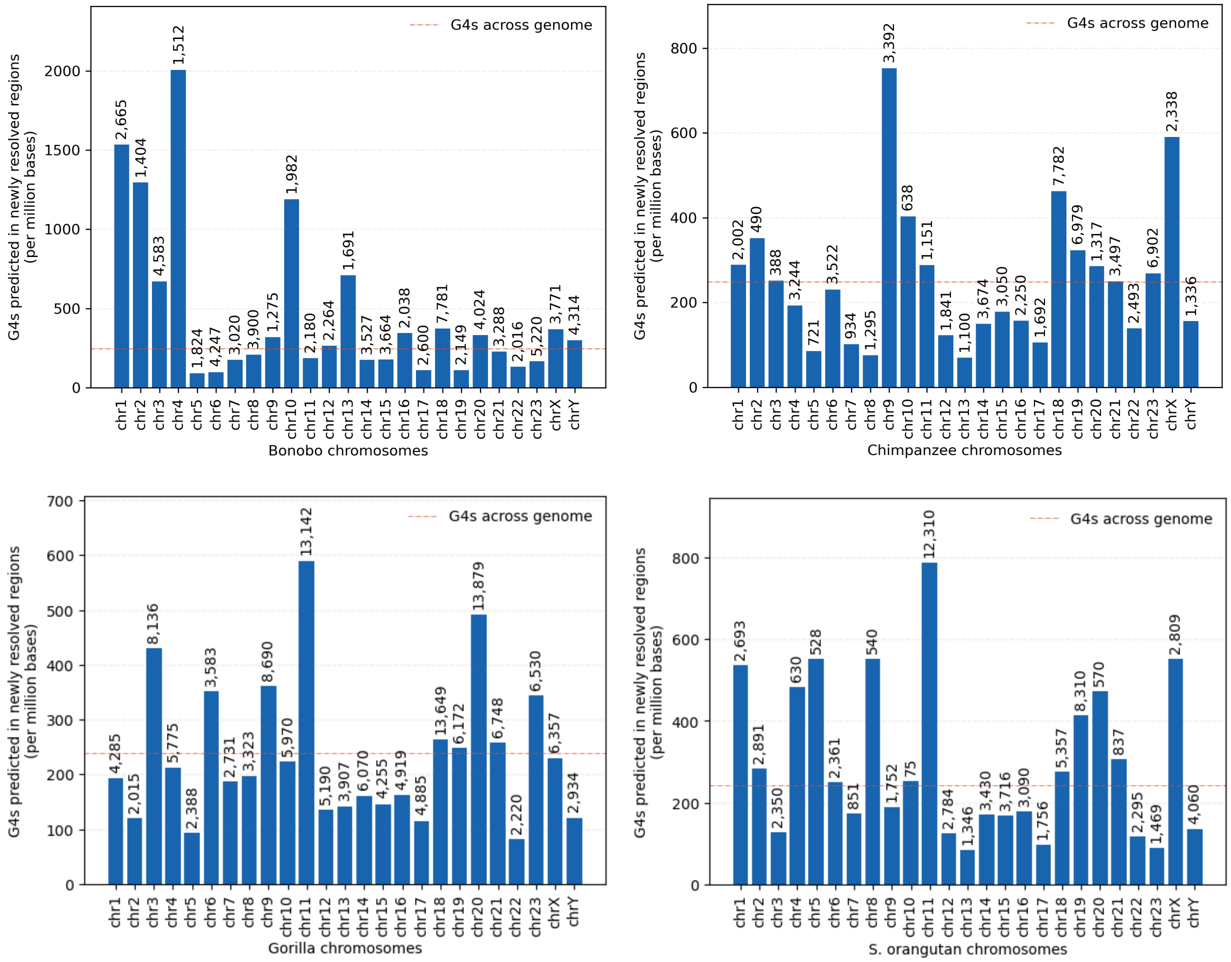

**Figure S3. Regression between G4 density and GC content of human chromosomes.**

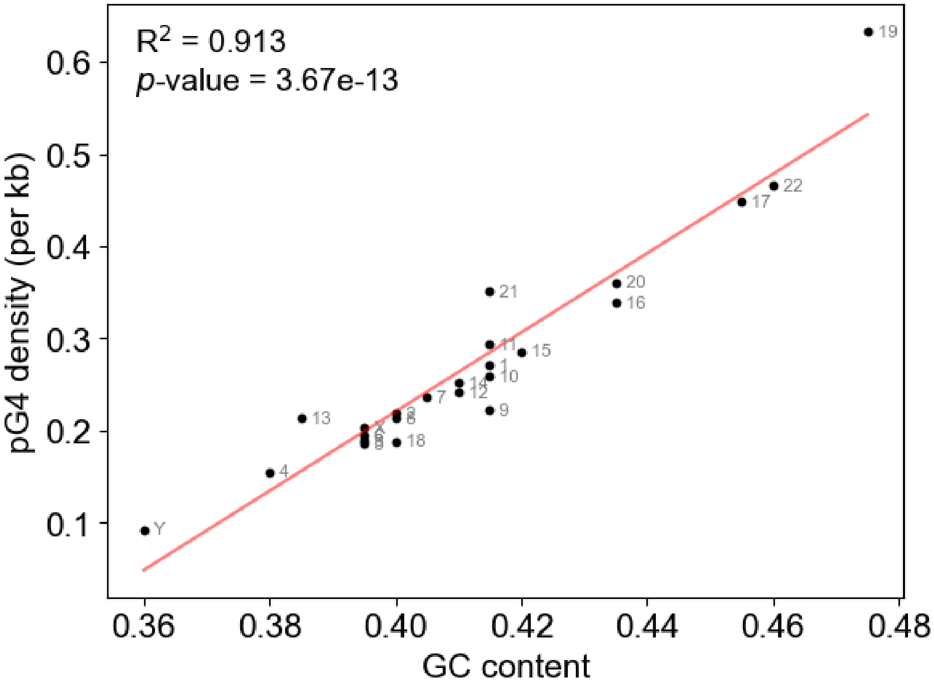

**Figure S4. Complete upset plots for the whole genome, HSA 1 through 22, X and Y.** The interspecific sharing of pG4s present at homologous locations of great ape T2T homologous chromosomes (HSA), 1 through 22, X, and Y. The vertical orange bars represent the number of pG4s shared across different great apes as shown by the shaded circles, and the vertical blue bars show species-specific G4s with aligned and unaligned G4s in dark and light blue, respectively. The horizontal bars represent the total number of G4s in each great ape species throughout its genome.

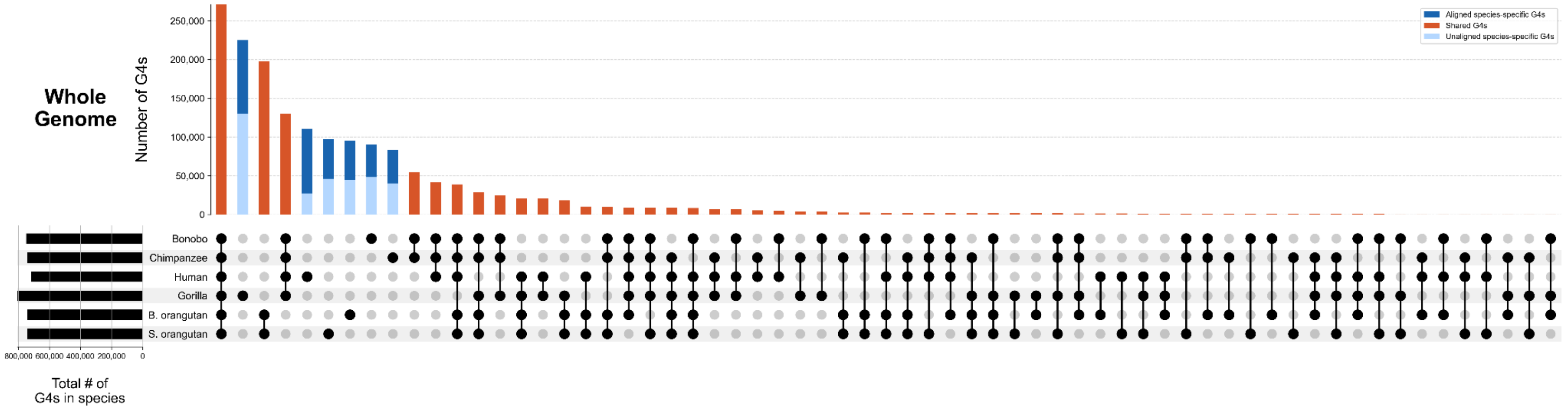

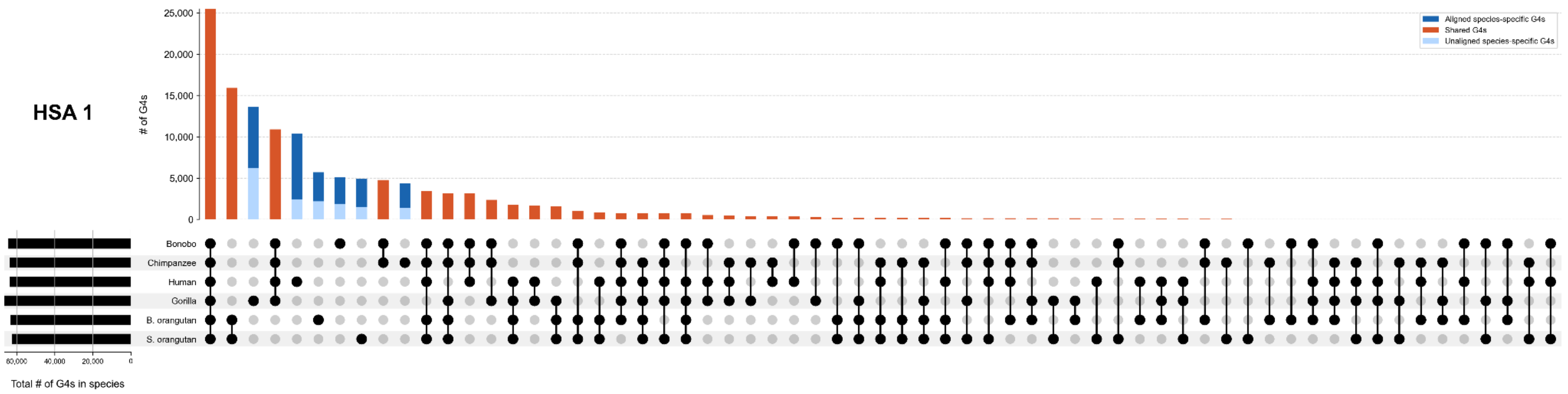
HSA 1
# of G4s
25,000
20,000
15,000
10,000
5,000
0
Aligned species-specific G4s
Shared G4s
Unaligned species-specific G4s
Bonobo
Chimpanzee
Human
Gorilla
B. orangutan
S. orangutan
60,000
40,000
20,000
0
Total # of G4s in species

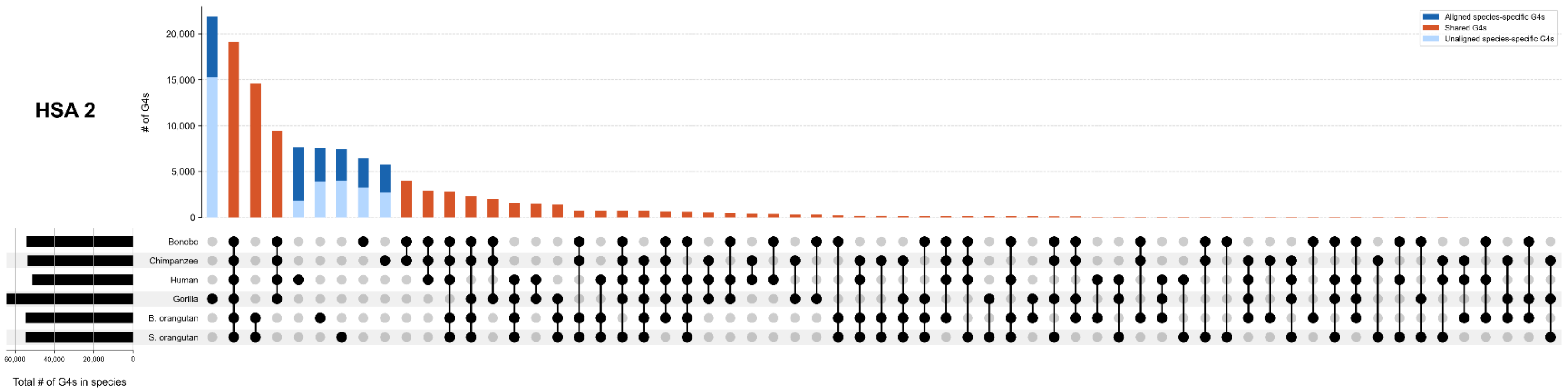
HSA 2
# of G4s
20,000
15,000
10,000
5,000
0
Aligned species-specific G4s
Shared G4s
Unaligned species-specific G4s
Bonobo
Chimpanzee
Human
Gorilla
B. orangutan
S. orangutan
60,000
40,000
20,000
0
Total # of G4s in species

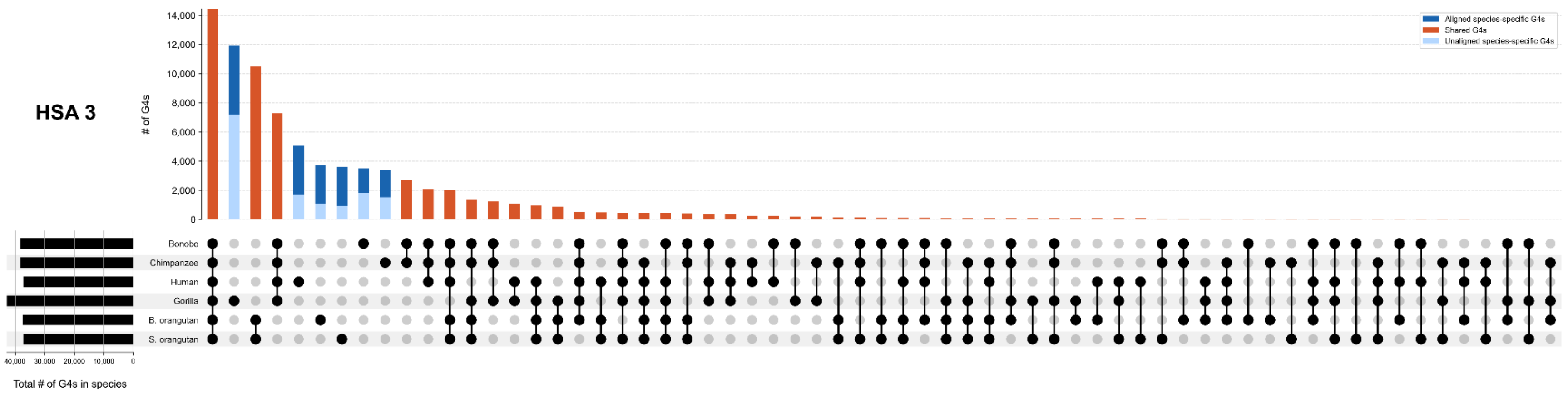
HSA 3
# of G4s
14,000
12,000
10,000
8,000
6,000
4,000
2,000
0
Aligned species-specific G4s
Shared G4s
Unaligned species-specific G4s
Bonobo
Chimpanzee
Human
Gorilla
B. orangutan
S. orangutan
40,000
30,000
20,000
10,000
0
Total # of G4s in species

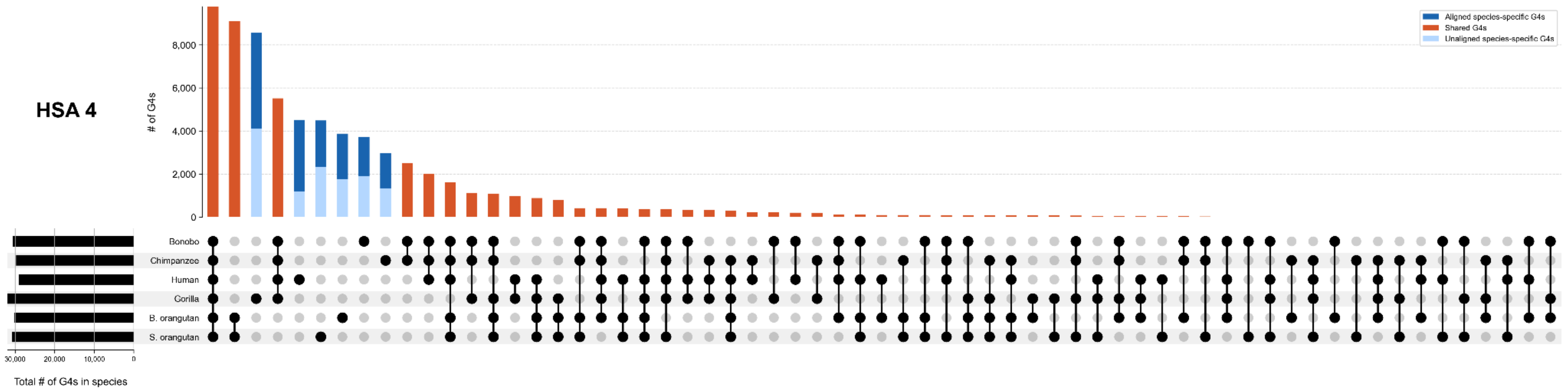
HSA 4
# of G4s
8,000
6,000
4,000
2,000
0
Aligned species-specific G4s
Shared G4s
Unaligned species-specific G4s
Bonobo
Chimpanzee
Human
Gorilla
B. orangutan
S. orangutan
30,000
20,000
10,000
0
Total # of G4s in species

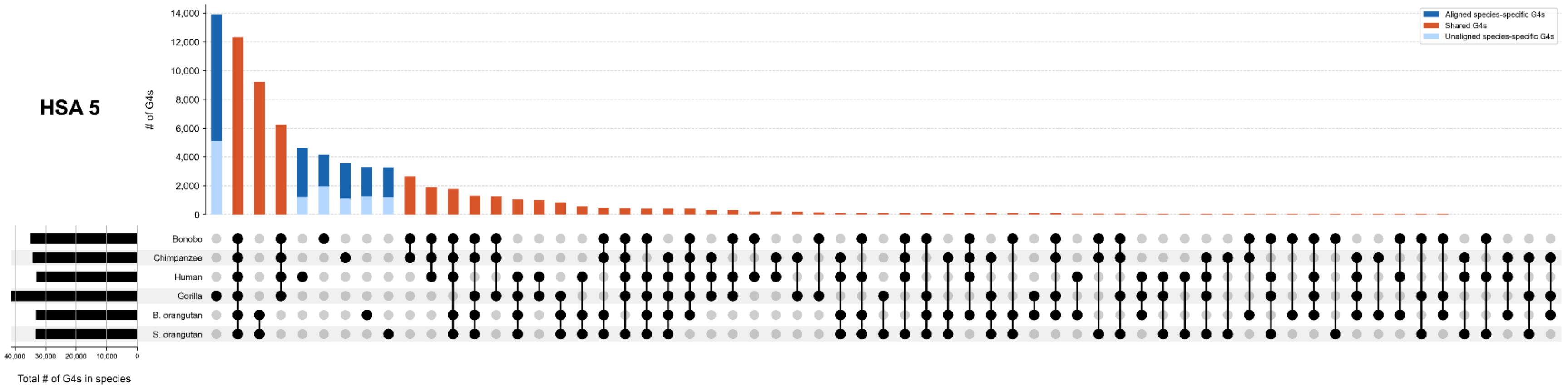
HSA 5
# of G4s
Aligned species-specific G4s
Shared G4s
Unaligned species-specific G4s
Bonobo
Chimpanzee
Human
Gorilla
B. orangutan
S. orangutan
Total # of G4s in species

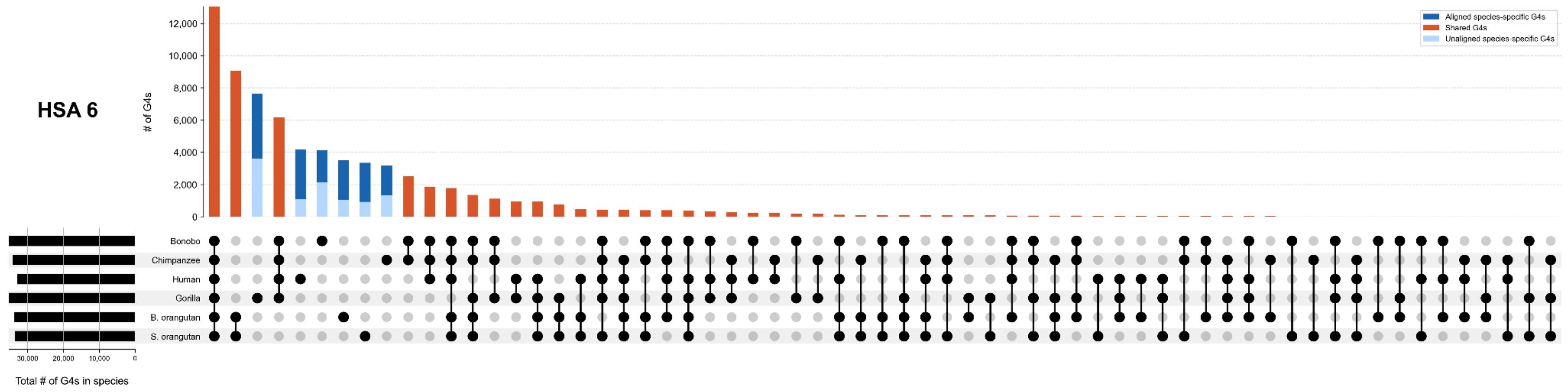
HSA 6
# of G4s
Aligned species-specific G4s
Shared G4s
Unaligned species-specific G4s
Bonobo
Chimpanzee
Human
Gorilla
B. orangutan
S. orangutan
Total # of G4s in species

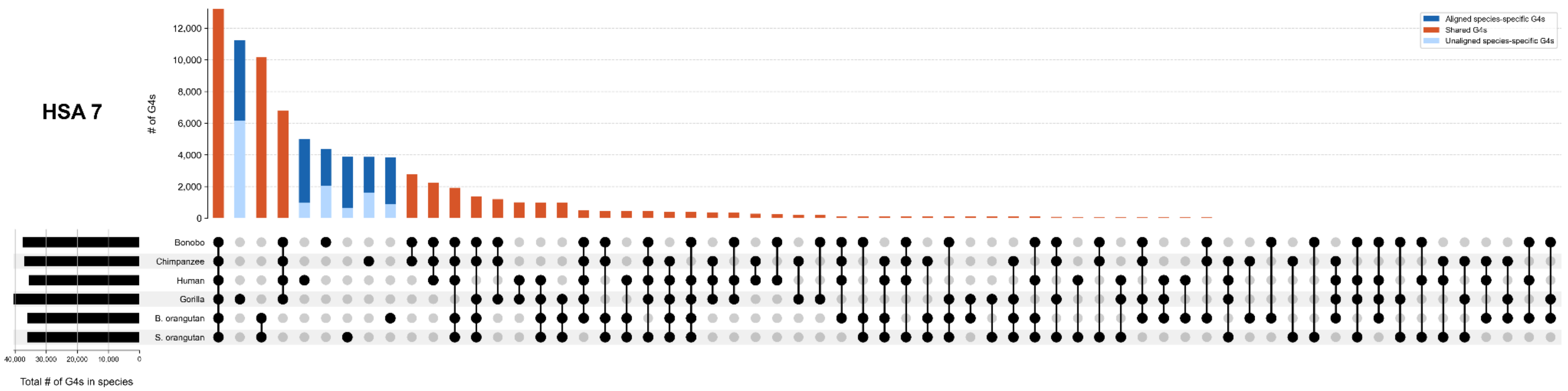

HSA 7
# of G4s
12,000
10,000
8,000
6,000
4,000
2,000
0
Aligned species-specific G4s
Shared G4s
Unaligned species-specific G4s
Bonobo
Chimpanzee
Human
Gorilla
B. orangutan
S. orangutan
40,000
30,000
20,000
10,000
0
Total # of G4s in species


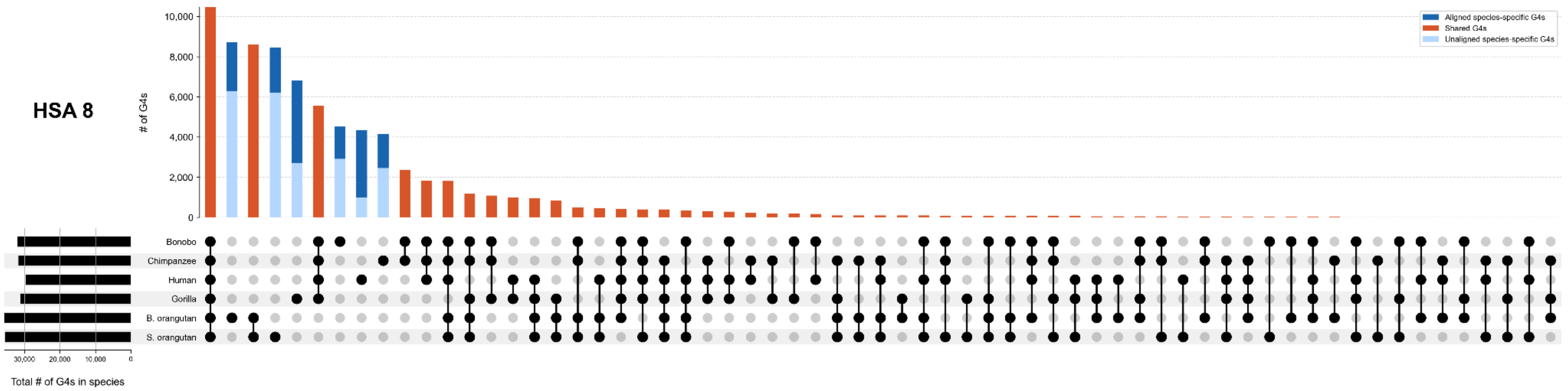

HSA 8
# of G4s
10,000
8,000
6,000
4,000
2,000
0
Aligned species-specific G4s
Shared G4s
Unaligned species-specific G4s
Bonobo
Chimpanzee
Human
Gorilla
B. orangutan
S. orangutan
30,000
20,000
10,000
0
Total # of G4s in species

HSA 9

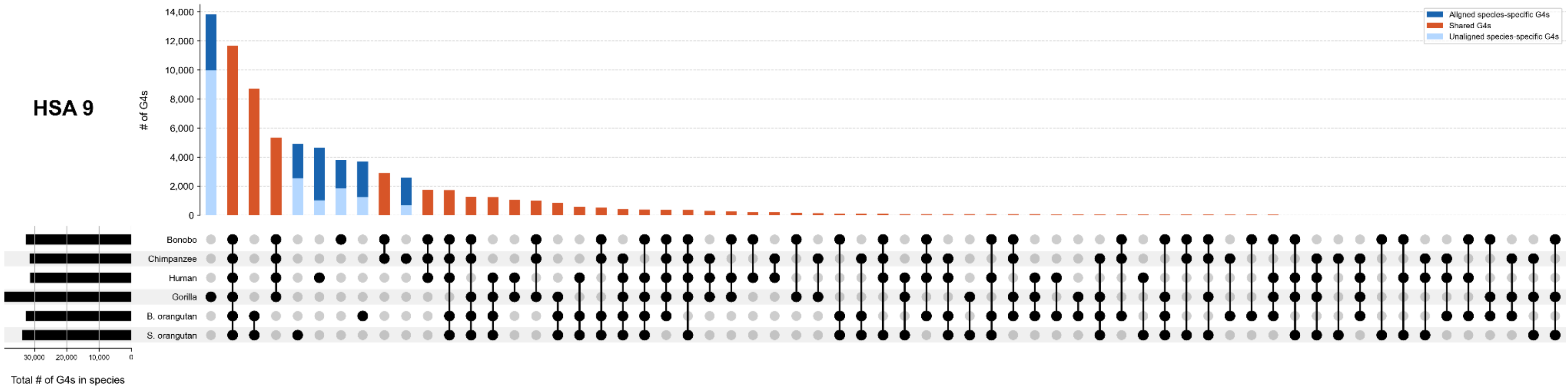

Aligned species-specific G4s
Shared G4s
Unaligned species-specific G4s
# of G4s
14,000
12,000
10,000
8,000
6,000
4,000
2,000
0
Bonobo
Chimpanzee
Human
Gorilla
B. orangutan
S. orangutan
30,000
20,000
10,000
0
Total # of G4s in species


HSA 10

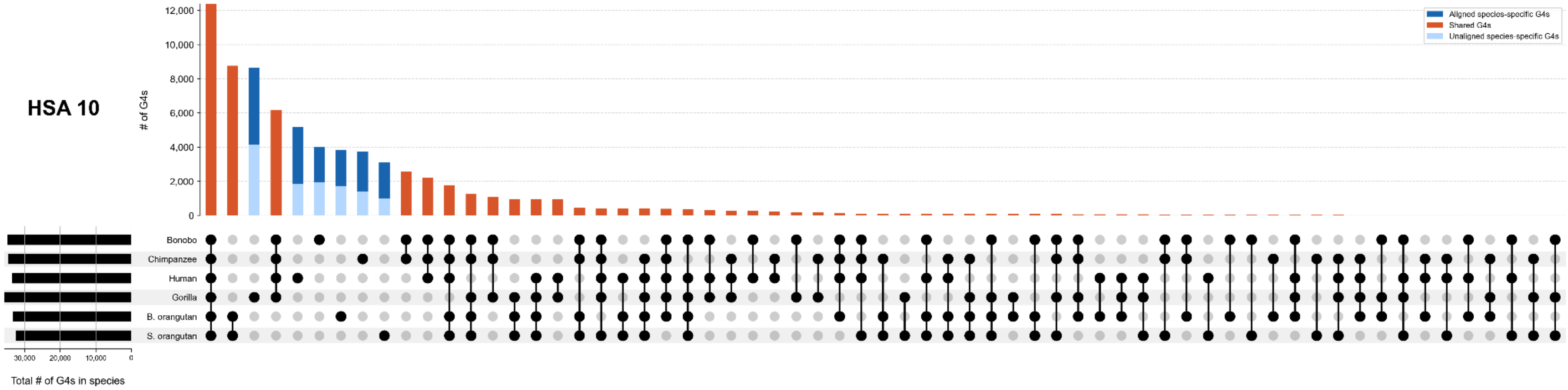

Aligned species-specific G4s
Shared G4s
Unaligned species-specific G4s
# of G4s
12,000
10,000
8,000
6,000
4,000
2,000
0
Bonobo
Chimpanzee
Human
Gorilla
B. orangutan
S. orangutan
30,000
20,000
10,000
0
Total # of G4s in species

HSA 11

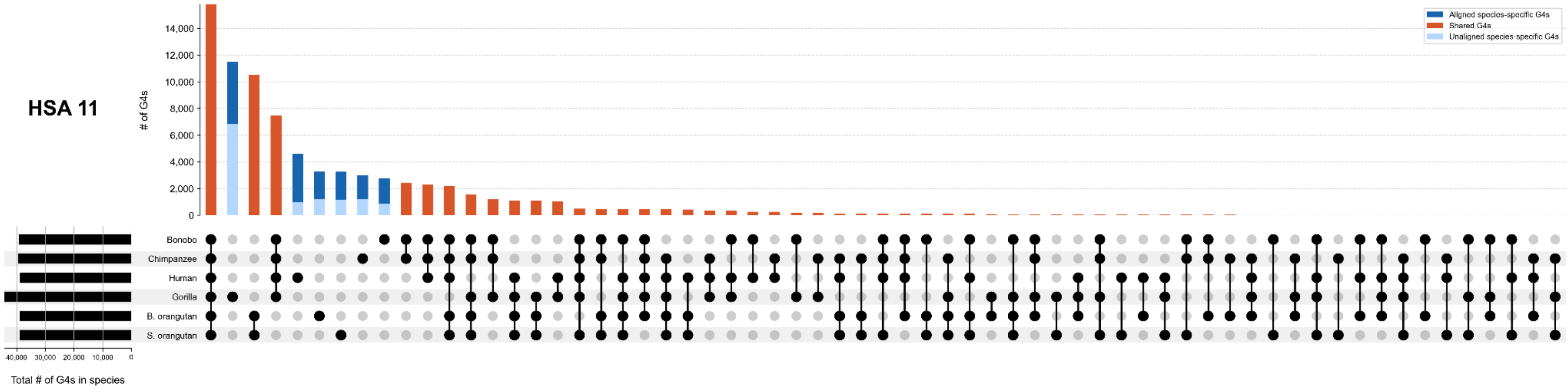

Aligned species-specific G4s
Shared G4s
Unaligned species-specific G4s
# of G4s
14,000
12,000
10,000
8,000
6,000
4,000
2,000
0
Bonobo
Chimpanzee
Human
Gorilla
B. orangutan
S. orangutan
40,000
30,000
20,000
10,000
0
Total # of G4s in species


HSA 12

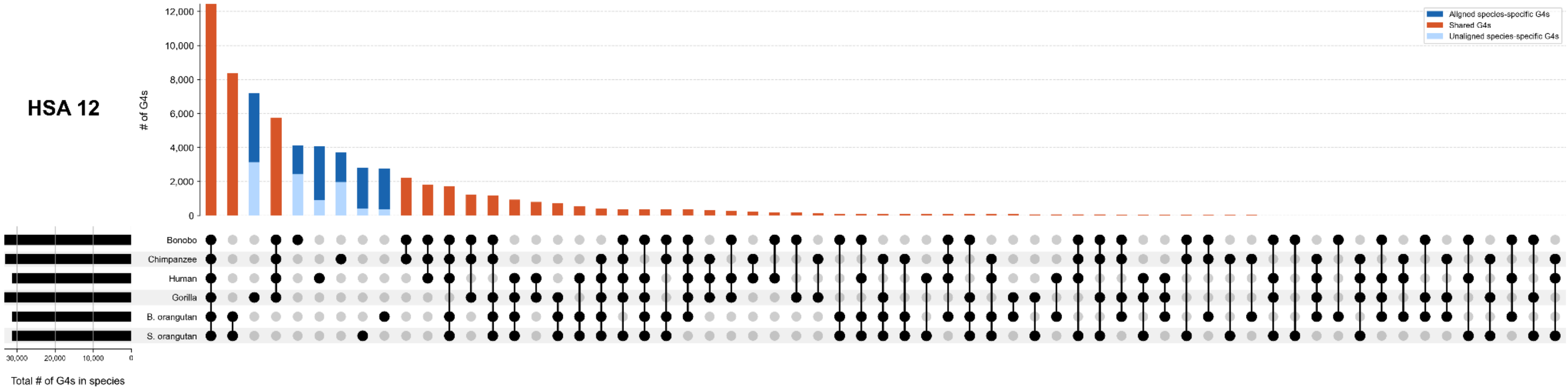

Aligned species-specific G4s
Shared G4s
Unaligned species-specific G4s
# of G4s
12,000
10,000
8,000
6,000
4,000
2,000
0
Bonobo
Chimpanzee
Human
Gorilla
B. orangutan
S. orangutan
30,000
20,000
10,000
0
Total # of G4s in species

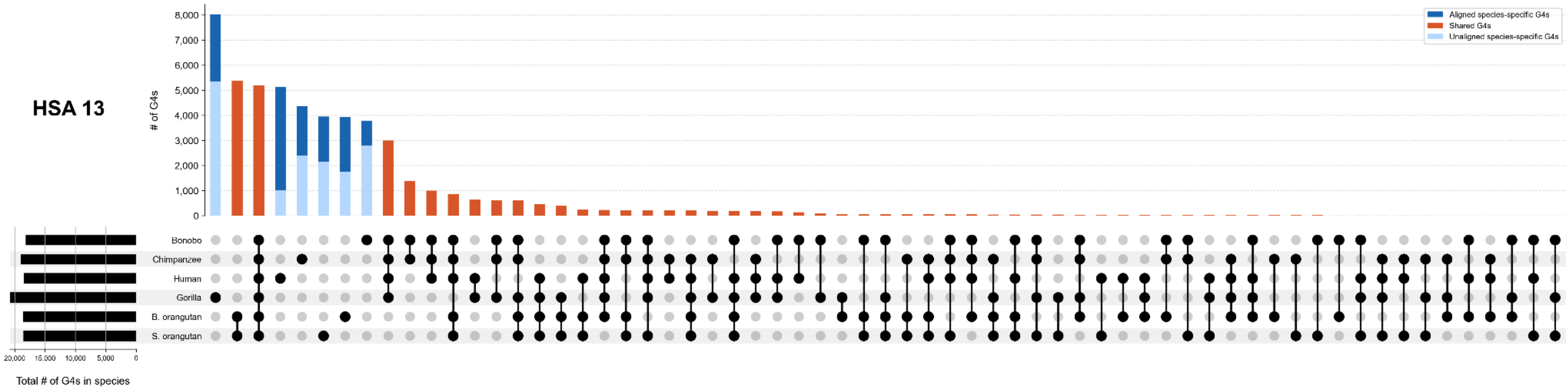
HSA 13
# of G4s
8,000
7,000
6,000
5,000
4,000
3,000
2,000
1,000
0
Aligned species-specific G4s
Shared G4s
Unaligned species-specific G4s
Bonobo
Chimpanzee
Human
Gorilla
B. orangutan
S. orangutan
20,000
15,000
10,000
5,000
0
Total # of G4s in species

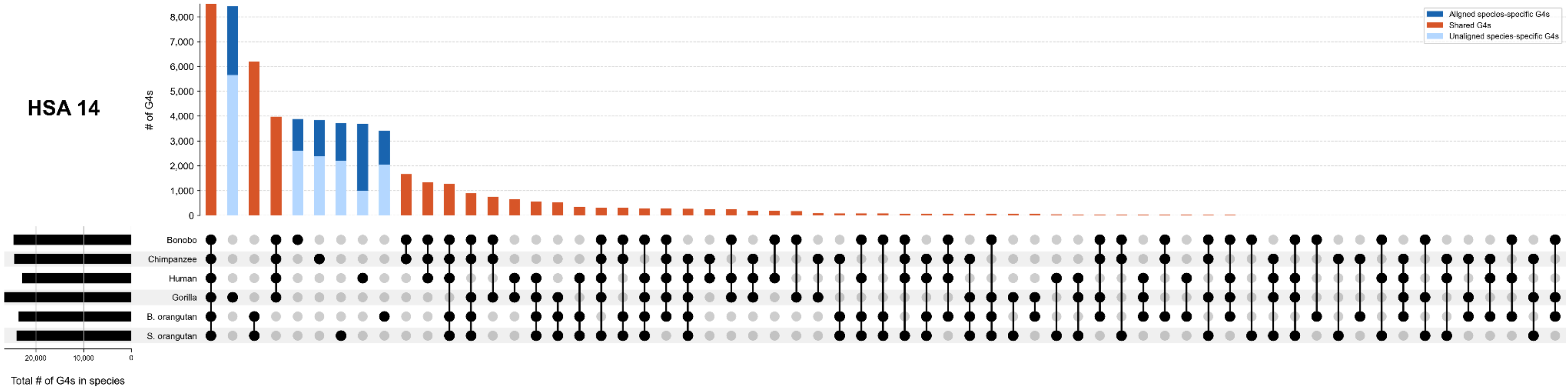
HSA 14
# of G4s
8,000
7,000
6,000
5,000
4,000
3,000
2,000
1,000
0
Aligned species-specific G4s
Shared G4s
Unaligned species-specific G4s
Bonobo
Chimpanzee
Human
Gorilla
B. orangutan
S. orangutan
20,000
10,000
0
Total # of G4s in species

#### HSA 15

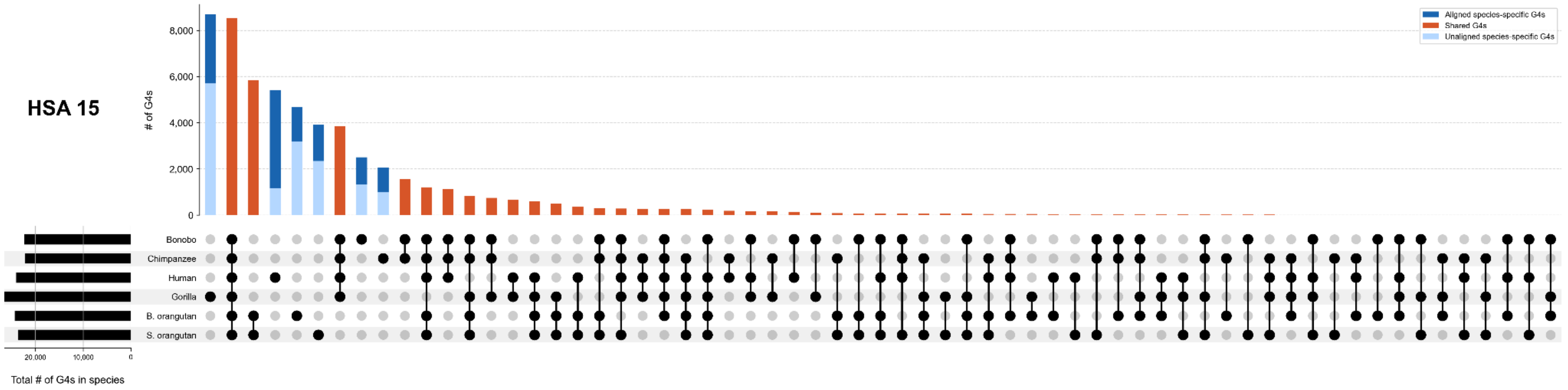

# of G4s
8,000
6,000
4,000
2,000
0
Aligned species-specific G4s
Shared G4s
Unaligned species-specific G4s
Bonobo
Chimpanzee
Human
Gorilla
B. orangutan
S. orangutan
20,000
10,000
0
Total # of G4s in species


#### HSA 16

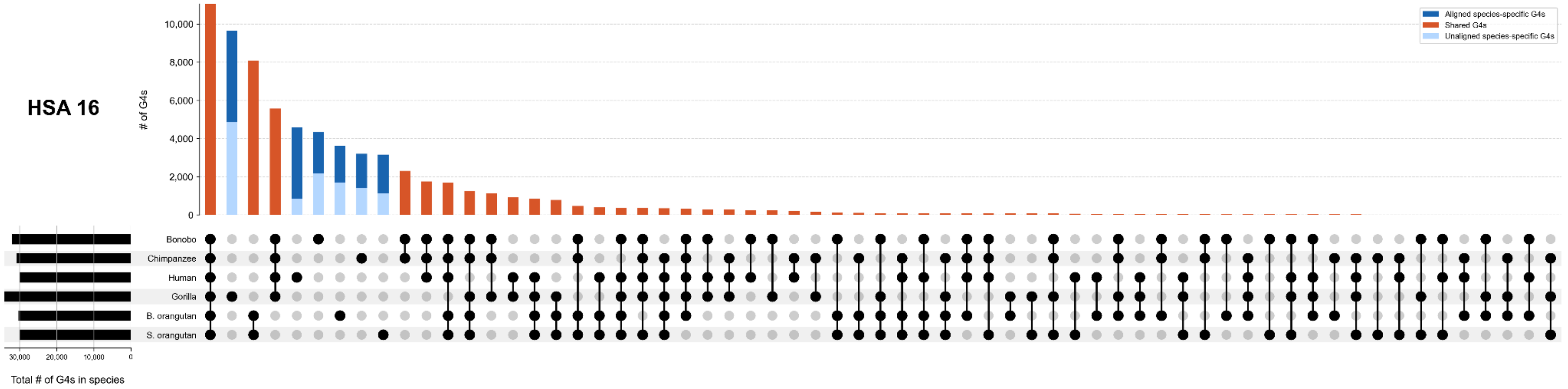

# of G4s
10,000
8,000
6,000
4,000
2,000
0
Aligned species-specific G4s
Shared G4s
Unaligned species-specific G4s
Bonobo
Chimpanzee
Human
Gorilla
B. orangutan
S. orangutan
30,000
20,000
10,000
0
Total # of G4s in species

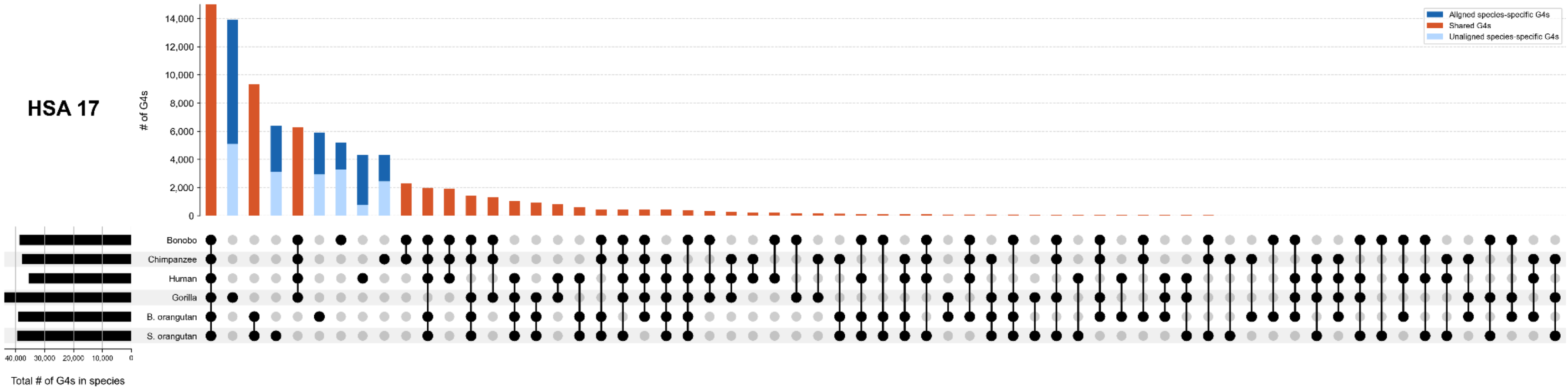

HSA 17
# of G4s
14,000
12,000
10,000
8,000
6,000
4,000
2,000
0
Aligned species-specific G4s
Shared G4s
Unaligned species-specific G4s
Bonobo
Chimpanzee
Human
Gorilla
B. orangutan
S. orangutan
40,000
30,000
20,000
10,000
0
Total # of G4s in species


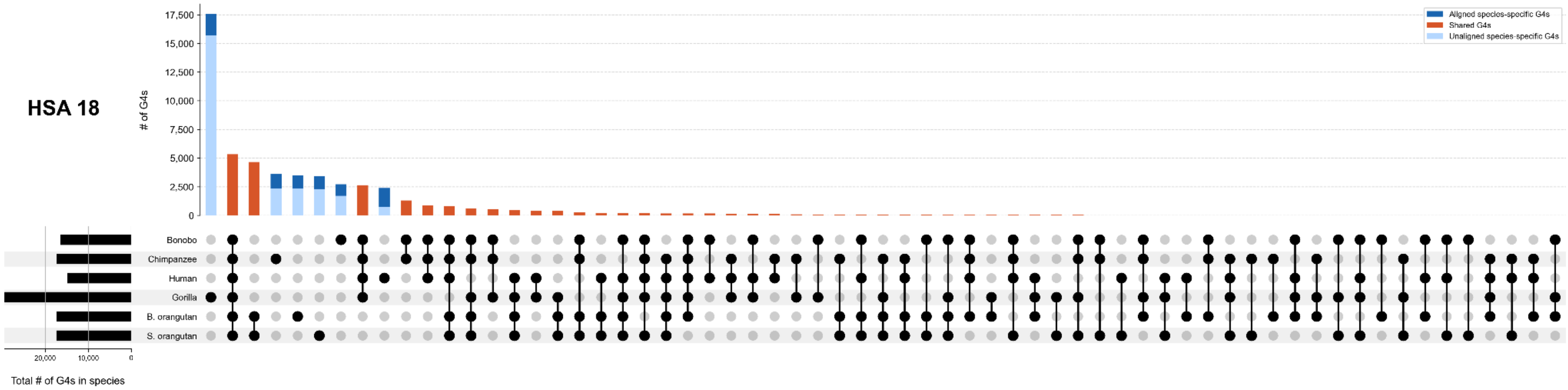

HSA 18
# of G4s
17,500
15,000
12,500
10,000
7,500
5,000
2,500
0
Aligned species-specific G4s
Shared G4s
Unaligned species-specific G4s
Bonobo
Chimpanzee
Human
Gorilla
B. orangutan
S. orangutan
20,000
10,000
0
Total # of G4s in species

HSA 19

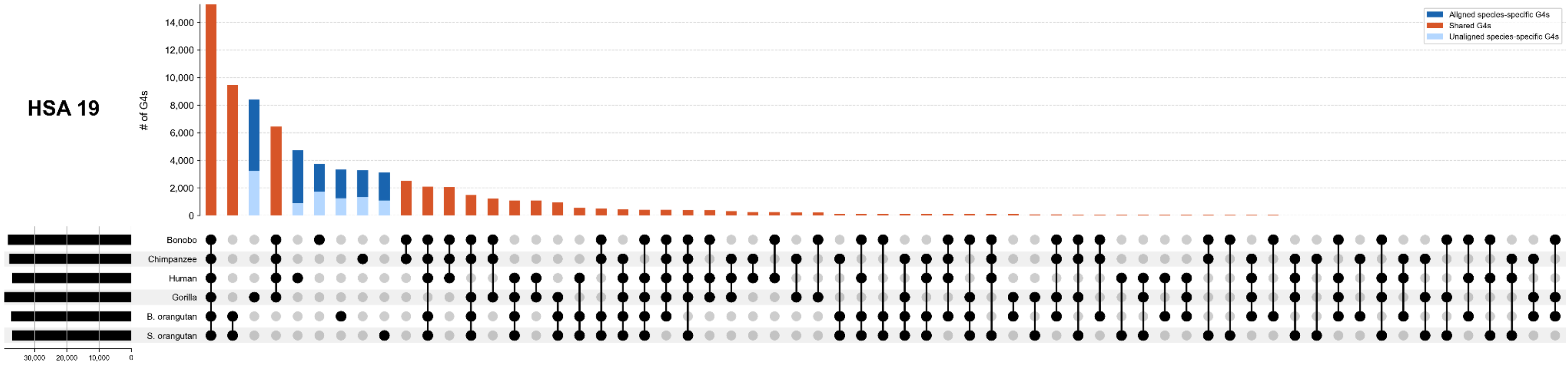
Aligned species-specific G4s
Shared G4s
Unaligned species-specific G4s
# of G4s
14,000
12,000
10,000
8,000
6,000
4,000
2,000
0
Bonobo
Chimpanzee
Human
Gorilla
B. orangutan
S. orangutan
30,000
20,000
10,000
0
Total # of G4s in species

HSA 20

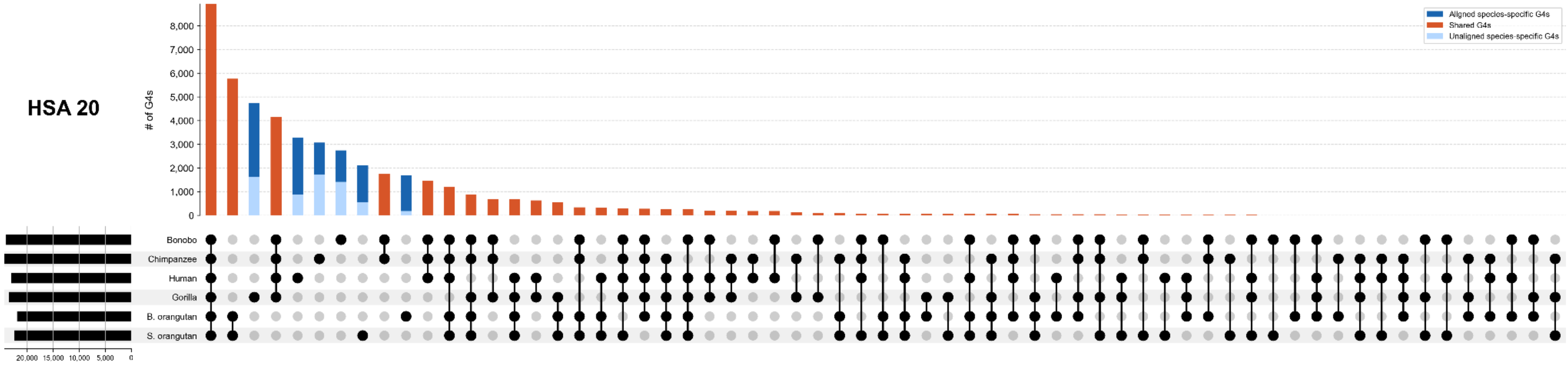
Aligned species-specific G4s
Shared G4s
Unaligned species-specific G4s
# of G4s
8,000
7,000
6,000
5,000
4,000
3,000
2,000
1,000
0
Bonobo
Chimpanzee
Human
Gorilla
B. orangutan
S. orangutan
20,000
15,000
10,000
5,000
0
Total # of G4s in species

HSA 21

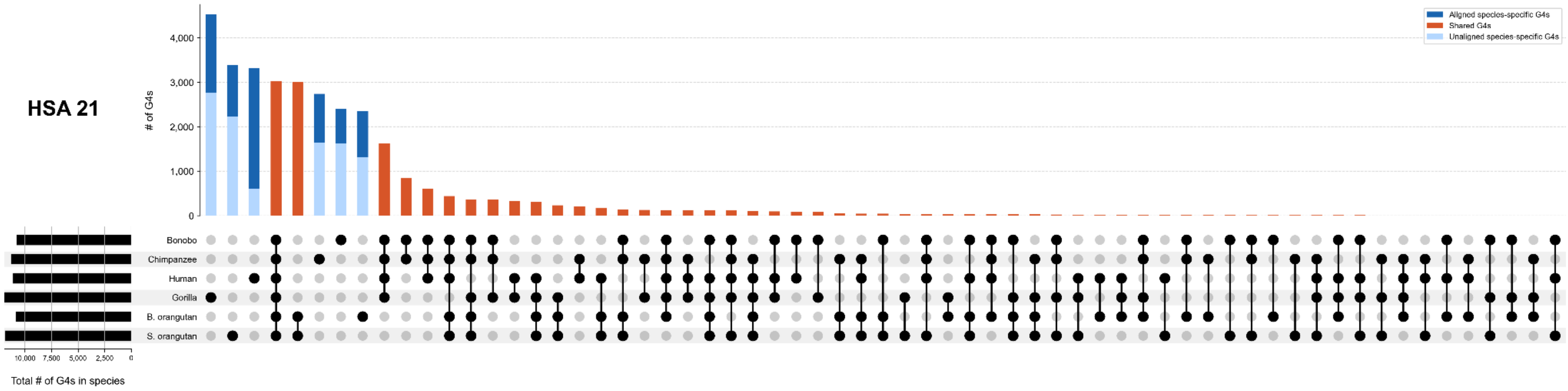

Aligned species-specific G4s
Shared G4s
Unaligned species-specific G4s
# of G4s
4,000
3,000
2,000
1,000
0
Bonobo
Chimpanzee
Human
Gorilla
B. orangutan
S. orangutan
10,000
7,500
5,000
2,500
0
Total # of G4s in species


HSA 22

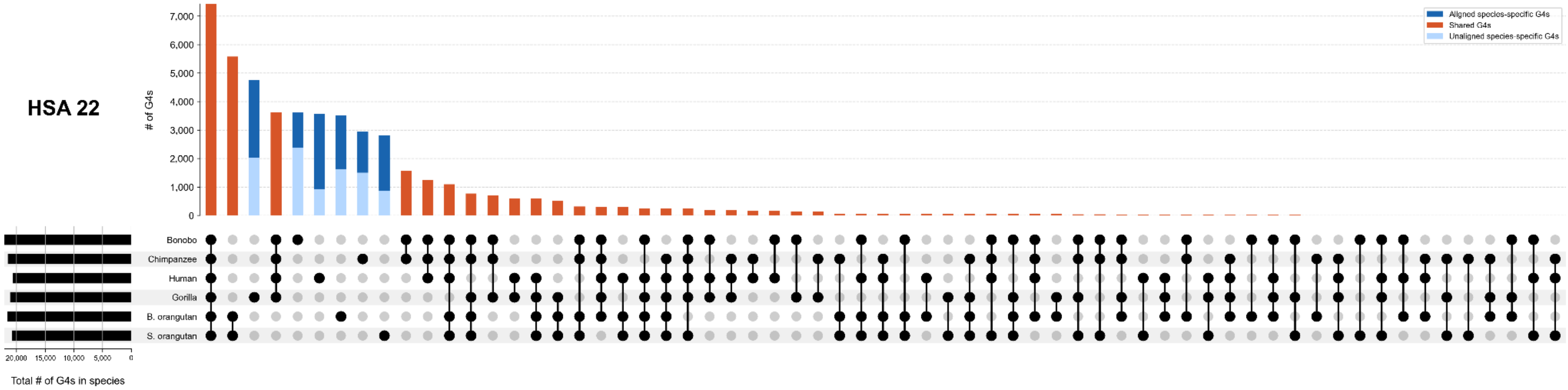

Aligned species-specific G4s
Shared G4s
Unaligned species-specific G4s
# of G4s
7,000
6,000
5,000
4,000
3,000
2,000
1,000
0
Bonobo
Chimpanzee
Human
Gorilla
B. orangutan
S. orangutan
20,000
15,000
10,000
5,000
0
Total # of G4s in species

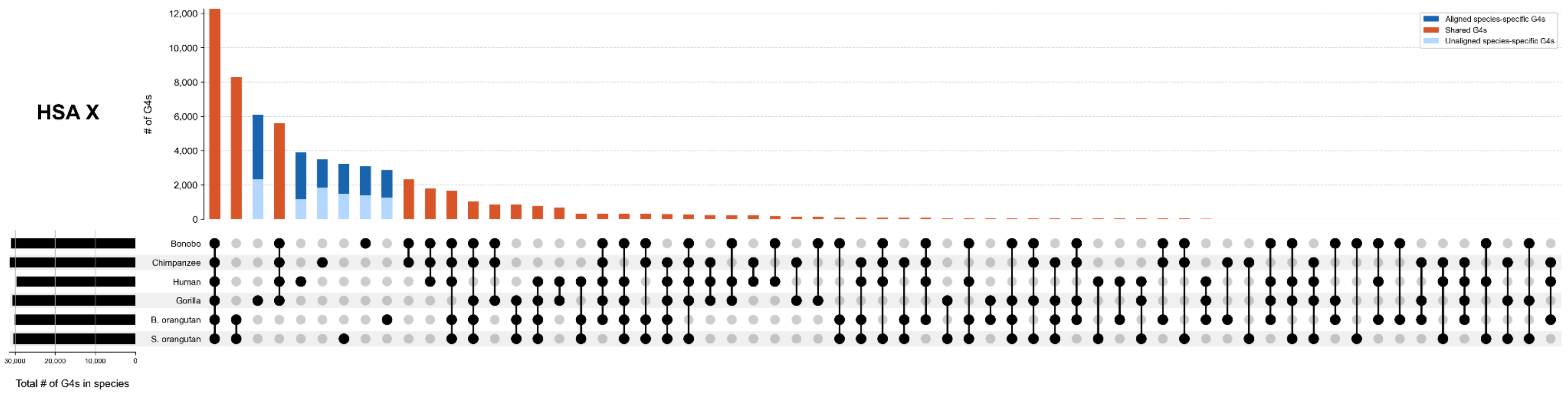
HSA X
# of G4s
12,000
10,000
8,000
6,000
4,000
2,000
0
Aligned species-specific G4s
Shared G4s
Unaligned species-specific G4s
Bonobo
Chimpanzee
Human
Gorilla
B. orangutan
S. orangutan
30,000
20,000
10,000
0
Total # of G4s in species

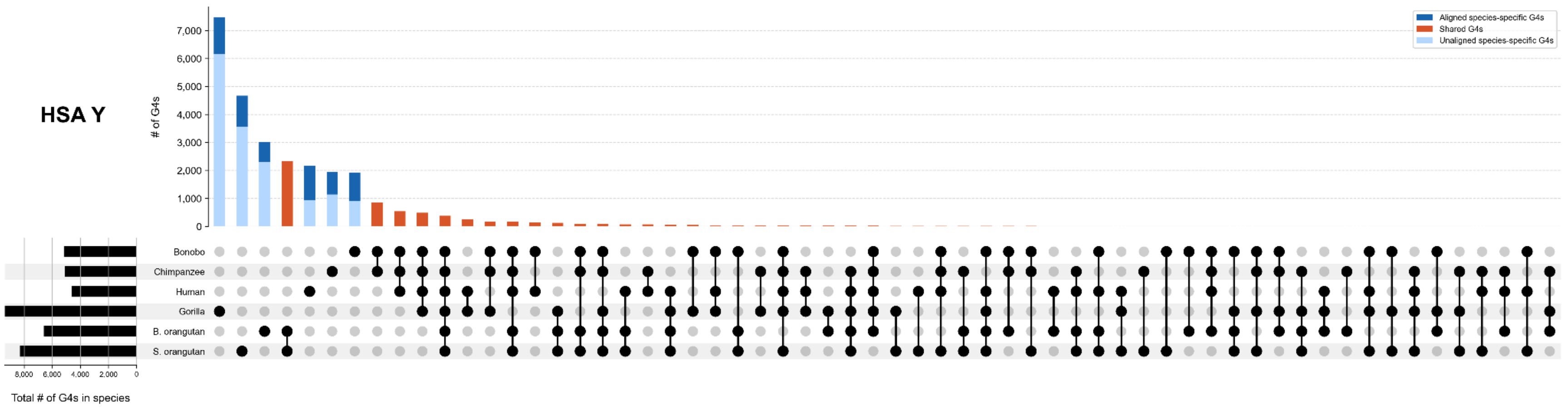
HSA Y
# of G4s
7,000
6,000
5,000
4,000
3,000
2,000
1,000
0
Aligned species-specific G4s
Shared G4s
Unaligned species-specific G4s
Bonobo
Chimpanzee
Human
Gorilla
B. orangutan
S. orangutan
8,000
6,000
4,000
2,000
0
Total # of G4s in species

**Figure S5. Species-specific pG4s across functional categories.** The number of aligned vs. unaligned species-specific G4s in each functional category across the genome—promoters, 5’UTRs, protein-coding sequences, introns, 3’UTRs, enhancers, non-protein-coding genes, origins of replication, CpG islands, repeats, and Non-Functional Non-Repetitive regions—across great apes.

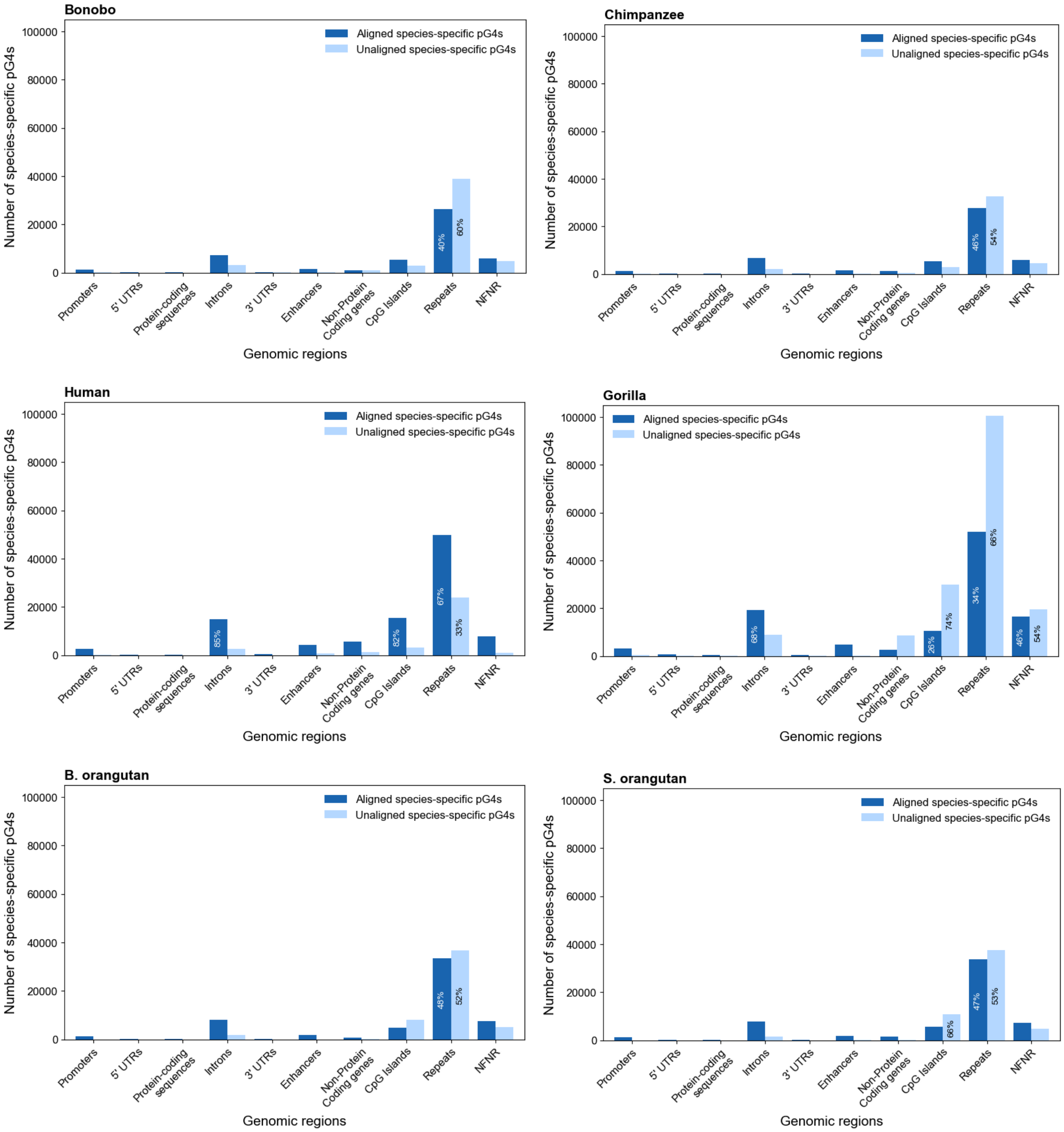

**Figure S6**. **Species-specific pG4s across repeat families genome-wide.** The number (and percentage) of aligned vs. unaligned species-specific G4s in repeat families (gorilla in Fig. 4C).

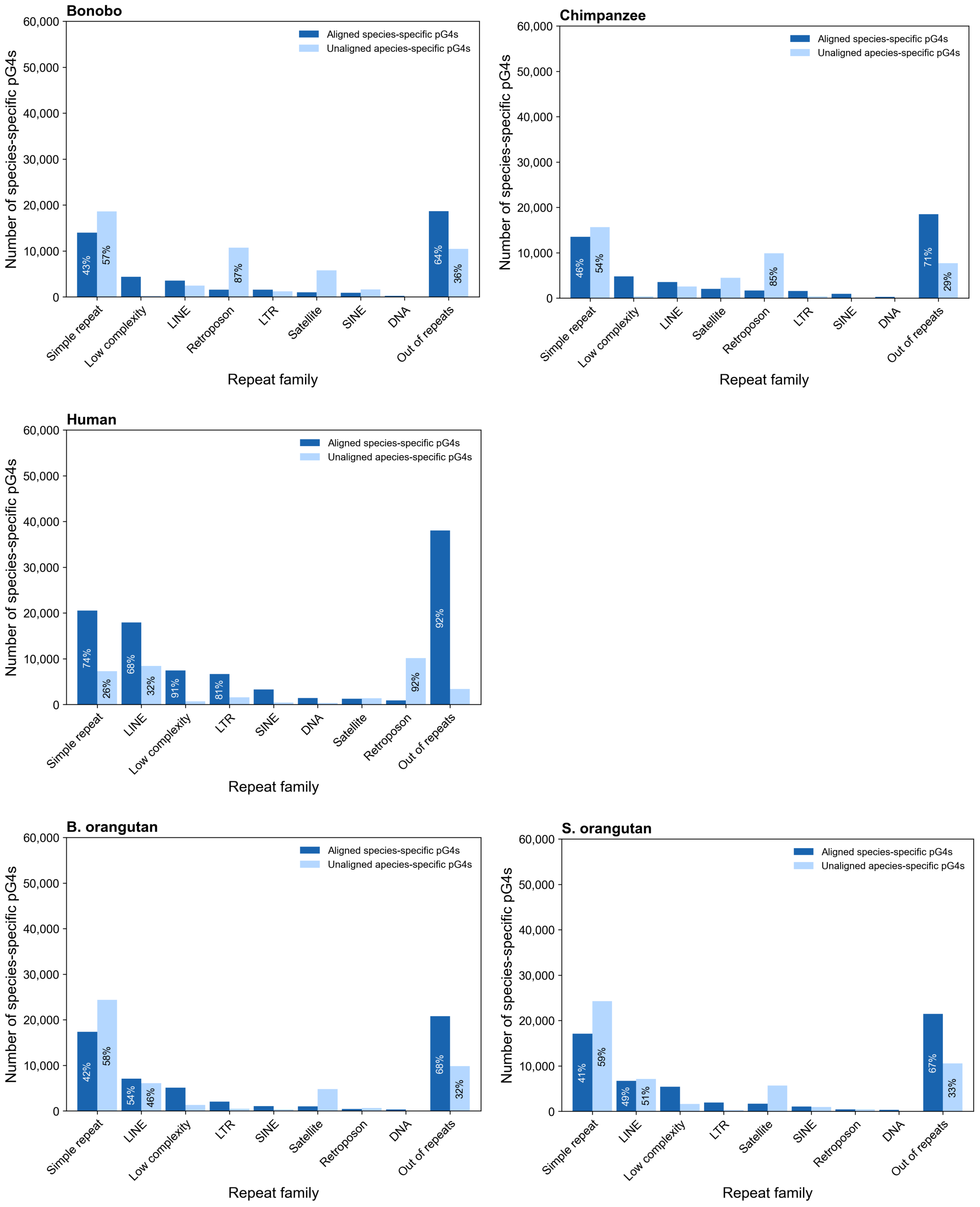

**Figure S7. Total number of non-shared G4s and divergence time, while excluding unaligned species-specific pG4s.** (A) The correlation between the total number of non-shared G4s, excluding the unaligned G4s, and divergence time in million years. Black dots represent pairs of species, and the red line represents the best fit. The correlation between the number of (B) aligned, and (C) unaligned species-specific G4s in great apes, and divergence times of each species in million years.

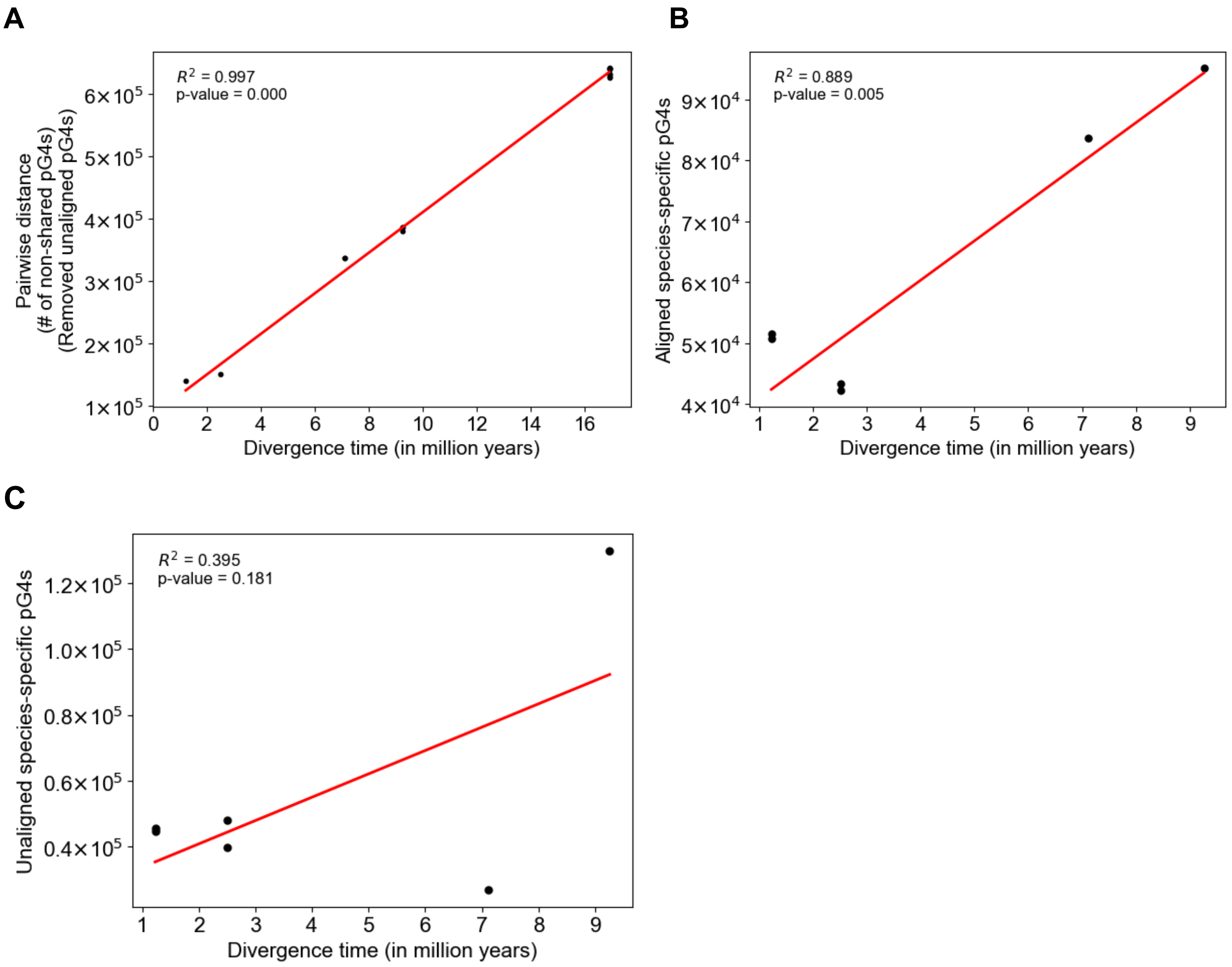

**Figure S8. pG4 enrichment and their methylation profiles at different functional categories.** The G4 enrichment, GC-corrected G4 enrichment, and methylation plots for different functional categories in (A-C) bonobo (fibroblast), (D-F) chimpanzee (lymphoblast), (G-I) gorilla (fibroblast), (J-L) B. orangutan (fibroblast), and (M-O) S. orangutan (fibroblast), respectively. The descriptions of the plot are identical to the ones described in Fig. 5A-C.

A

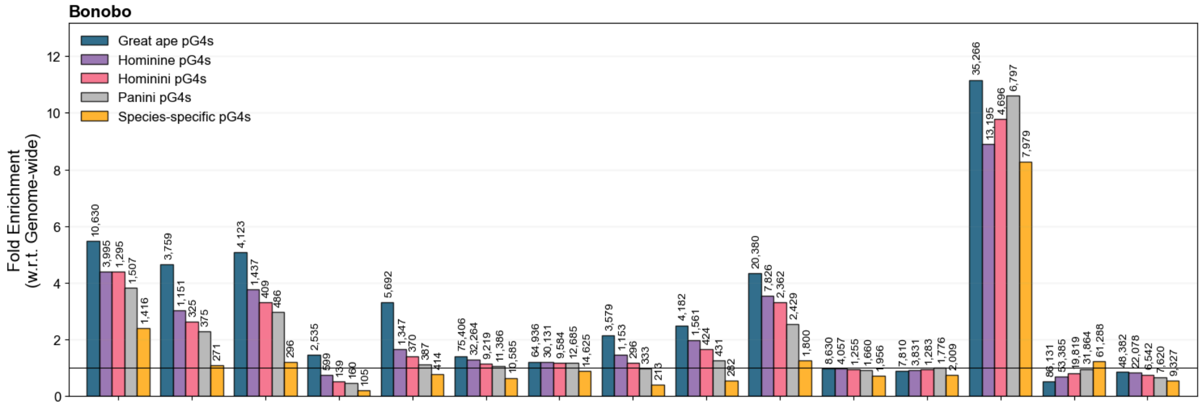


B

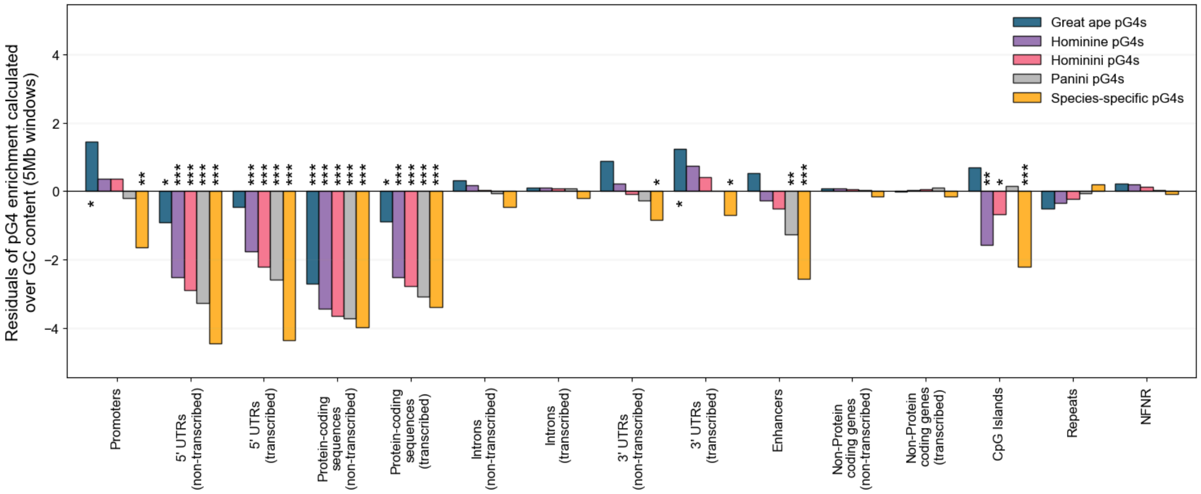


C

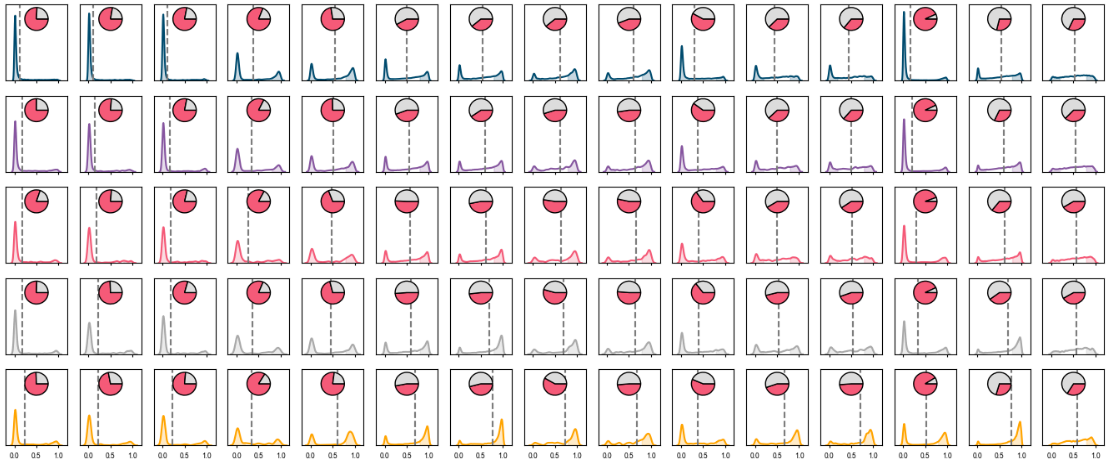

**D**

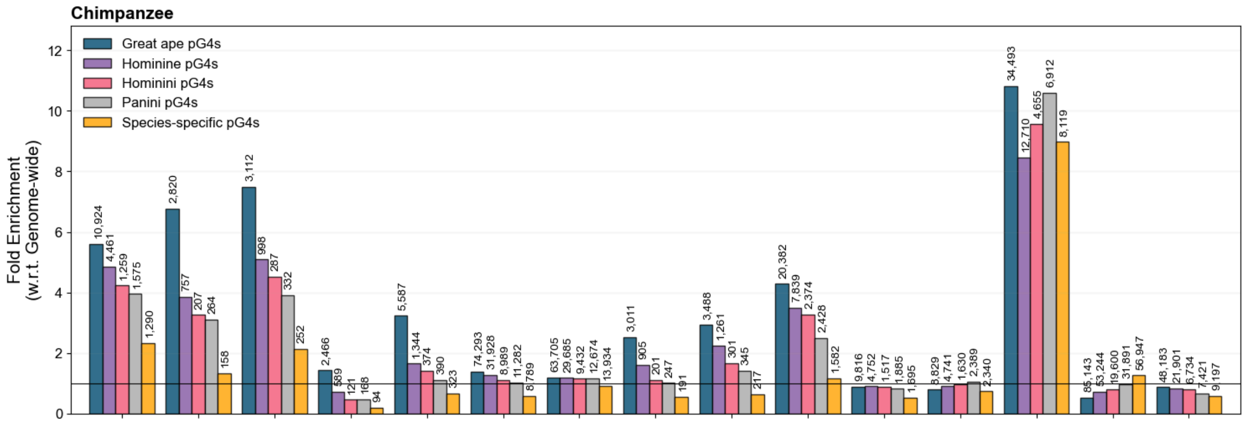
Chimpanzee
Great ape pG4s
Hominine pG4s
Hominini pG4s
Panini pG4s
Species-specific pG4s
Fold Enrichment
(w.r.t. Genome-wide)

**E**

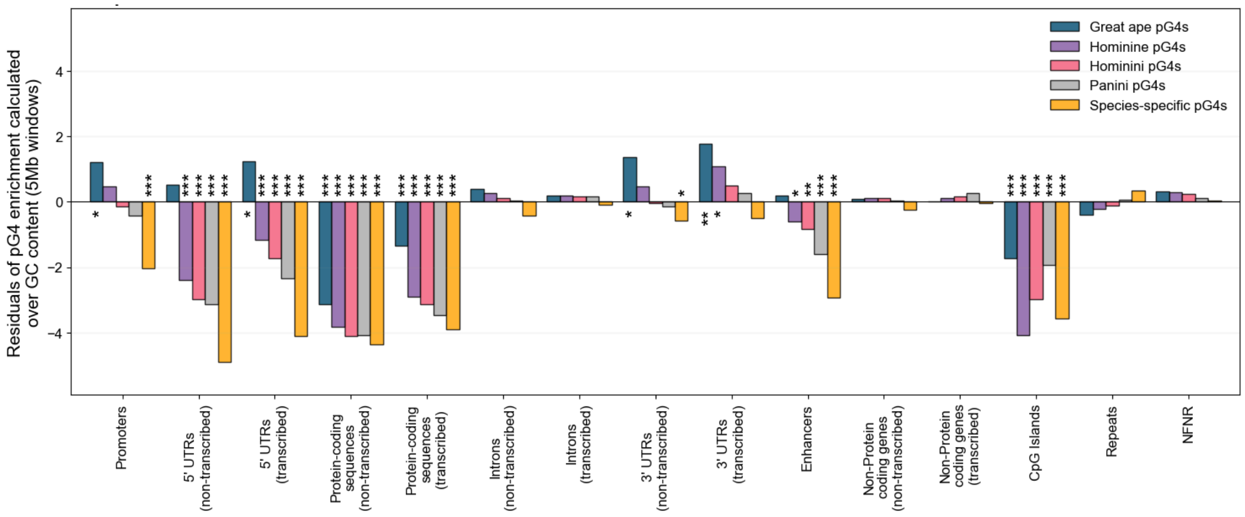
Great ape pG4s
Hominine pG4s
Hominini pG4s
Panini pG4s
Species-specific pG4s
Residuals of pG4 enrichment calculated
over GC content (5Mb windows)
Promoters
5' UTRs
(non-transcribed)
5' UTRs
(transcribed)
Protein-coding
sequences
(non-transcribed)
Protein-coding
sequences
(transcribed)
Introns
(non-transcribed)
Introns
(transcribed)
3' UTRs
(non-transcribed)
3' UTRs
(transcribed)
Enhancers
Non-Protein
coding genes
(non-transcribed)
Non-Protein
coding genes
(transcribed)
CpG Islands
Repeats
NFNR

**F**

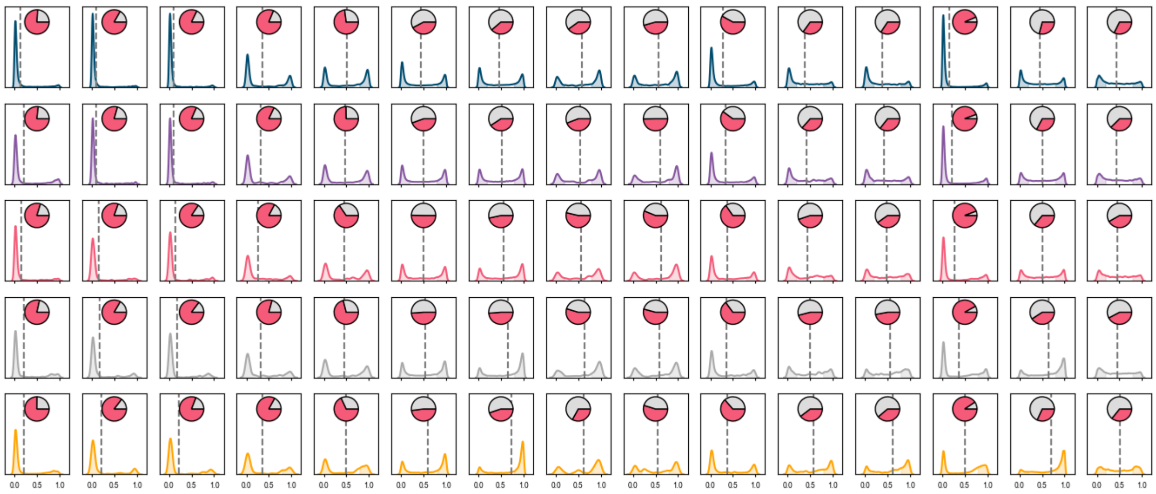

**G**

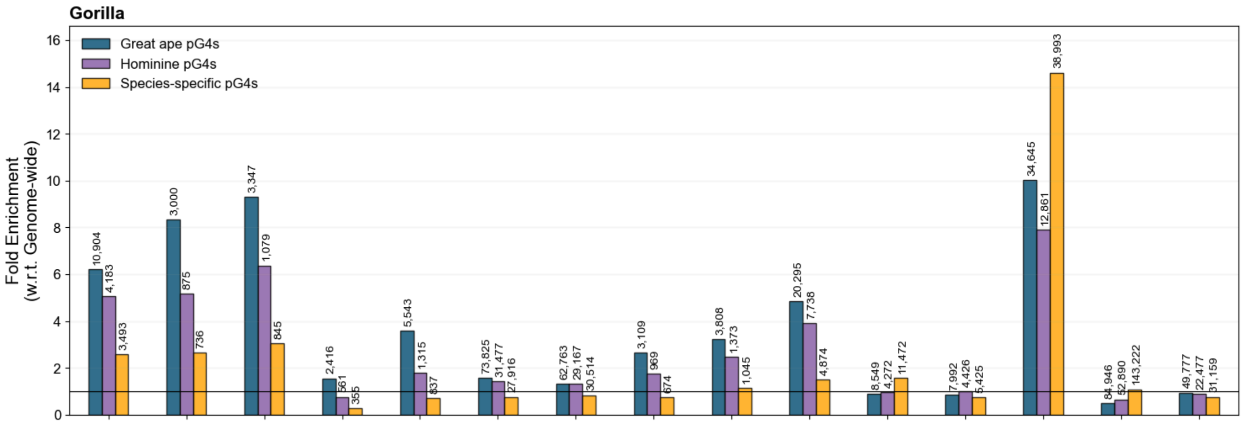
Gorilla
Great ape pG4s
Hominine pG4s
Species-specific pG4s
Fold Enrichment (w.r.t. Genome-wide)
0
2
4
6
8
10
12
14
16
10,904 4,183 3,493
3,000 875 736
3,347 1,079 845
2,416 561 355
5,543 1,315 837
73,825 31,477 27,916
62,763 29,167 30,514
3,109 969 674
3,808 1,373 1,045
20,295 7,738 4,874
8,549 4,272 11,472
7,992 4,426 5,425
34,645 12,861 38,993
84,946 52,890 143,222
49,777 22,477 31,159

**H**

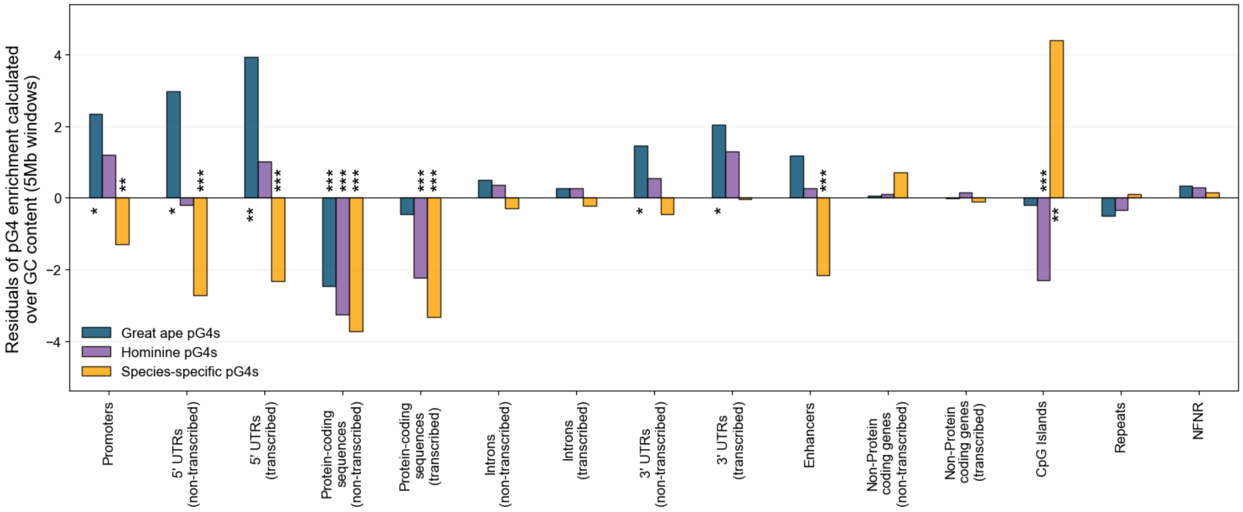
Residuals of pG4 enrichment calculated over GC content (5Mb windows)
-4
-2
0
2
4
Great ape pG4s
Hominine pG4s
Species-specific pG4s
Promoters
5' UTRs (non-transcribed)
5' UTRs (transcribed)
Protein-coding sequences (non-transcribed)
Protein-coding sequences (transcribed)
Introns (non-transcribed)
Introns (transcribed)
3' UTRs (non-transcribed)
3' UTRs (transcribed)
Enhancers
Non-Protein coding genes (non-transcribed)
Non-Protein coding genes (transcribed)
CpG Islands
Repeats
NFNR

**I**

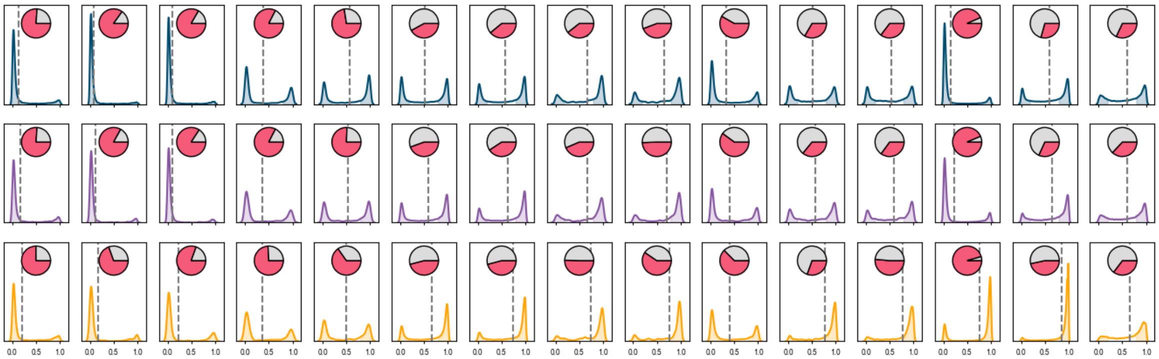
0.0 0.5 1.0

J

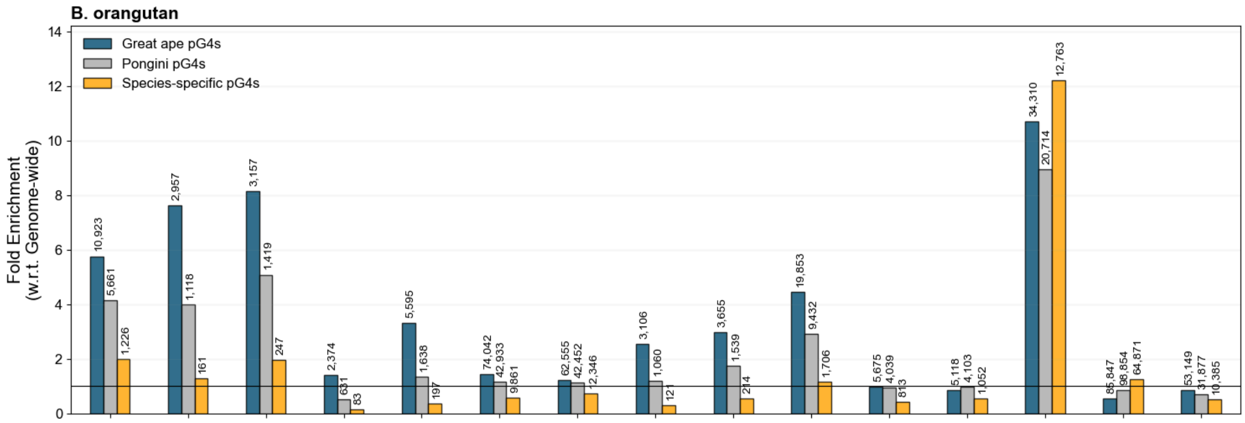

B. orangutan
Fold Enrichment (w.r.t. Genome-wide)
Great ape pG4s
Pongini pG4s
Species-specific pG4s
10,923
5,661
1,226
2,957
1,118
161
3,157
1,419
247
2,374
631
83
5,595
1,638
197
74,042
42,933
9,861
62,555
42,452
2,346
3,106
1,060
121
3,655
1,539
214
19,853
9,432
1,706
5,675
4,039
813
5,118
4,103
1,052
34,310
20,714
12,763
85,847
98,854
64,871
53,149
31,877
10,385


K

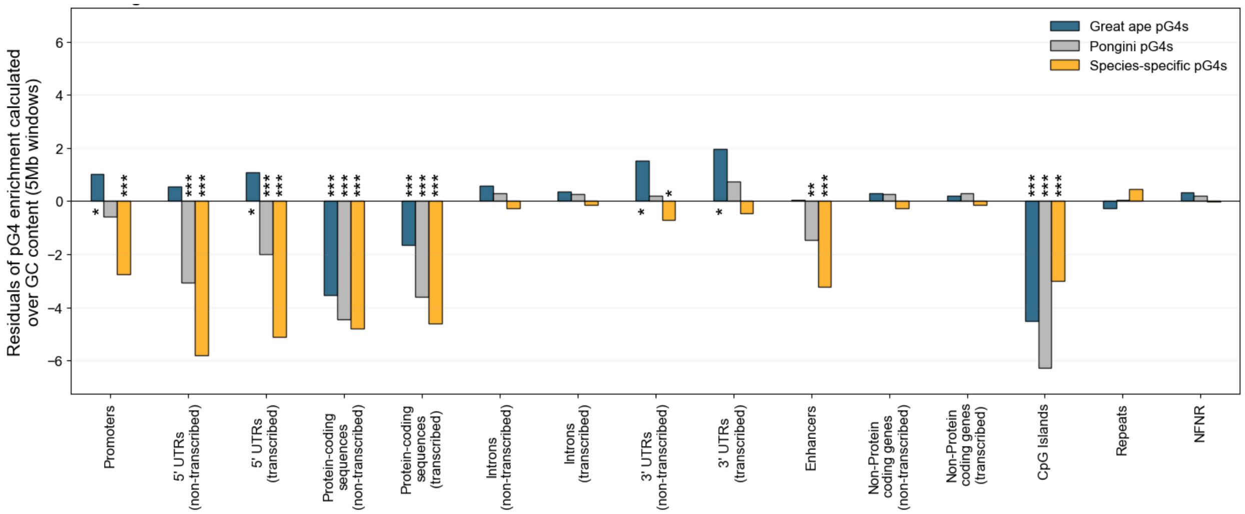

Residuals of pG4 enrichment calculated over GC content (5Mb windows)
Great ape pG4s
Pongini pG4s
Species-specific pG4s
Promoters
5' UTRs (non-transcribed)
5' UTRs (transcribed)
Protein-coding sequences (non-transcribed)
Protein-coding sequences (transcribed)
Introns (non-transcribed)
Introns (transcribed)
3' UTRs (non-transcribed)
3' UTRs (transcribed)
Enhancers
Non-Protein coding genes (non-transcribed)
Non-Protein coding genes (transcribed)
CpG Islands
Repeats
NFNR


L

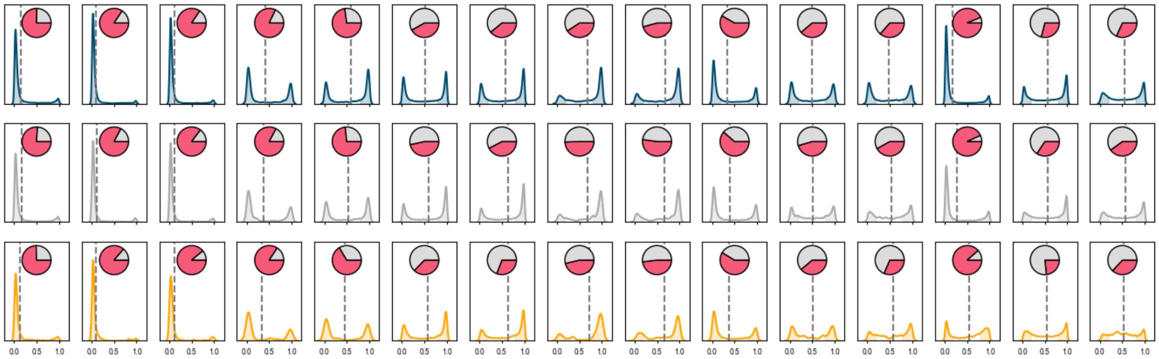

**M**

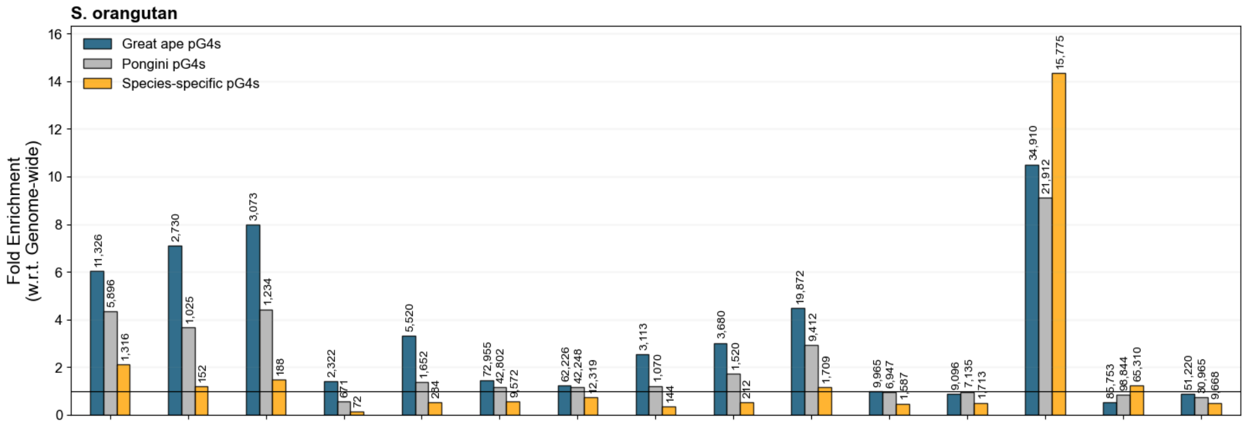
S. orangutan
Fold Enrichment (w.r.t. Genome-wide)
Great ape pG4s
Pongini pG4s
Species-specific pG4s
11,326
5,896
1,316
2,730
1,025
152
3,073
1,234
188
2,322
671
72
5,520
1,652
284
72,955
42,802
9,572
62,226
42,248
12,319
3,113
1,070
144
3,680
1,520
212
19,872
9,412
1,709
9,965
6,947
1,587
9,096
7,135
1,713
34,910
21,912
15,775
85,753
98,844
65,310
51,220
30,965
9,668

**N**

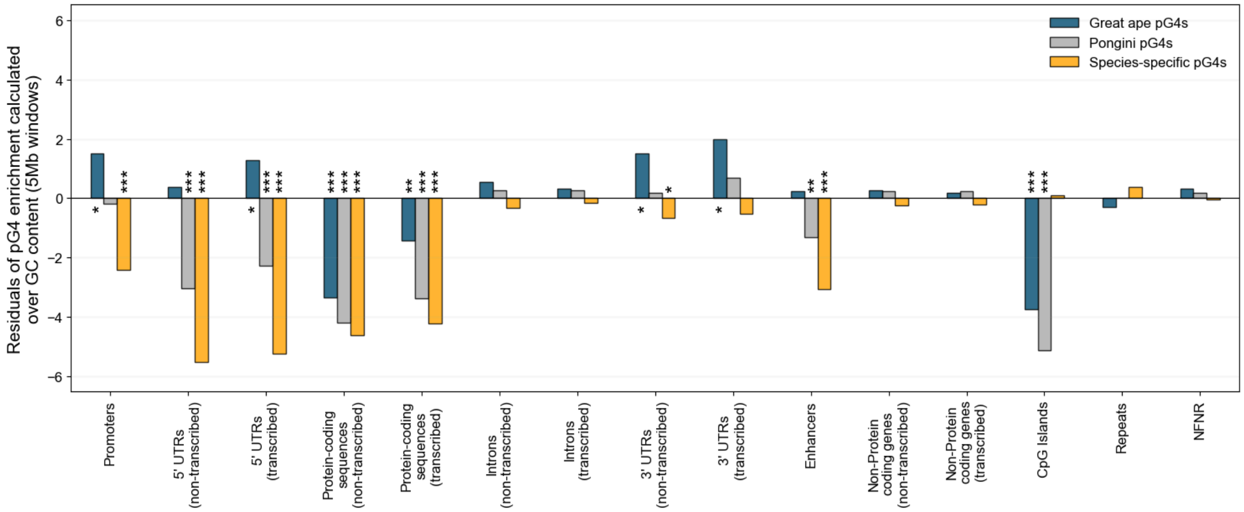
Residuals of pG4 enrichment calculated over GC content (5Mb windows)
Great ape pG4s
Pongini pG4s
Species-specific pG4s
Promoters
5' UTRs (non-transcribed)
5' UTRs (transcribed)
Protein-coding sequences (non-transcribed)
Protein-coding sequences (transcribed)
Introns (non-transcribed)
Introns (transcribed)
3' UTRs (non-transcribed)
3' UTRs (transcribed)
Enhancers
Non-Protein coding genes (non-transcribed)
Non-Protein coding genes (transcribed)
CpG Islands
Repeats
NFNR

**O**

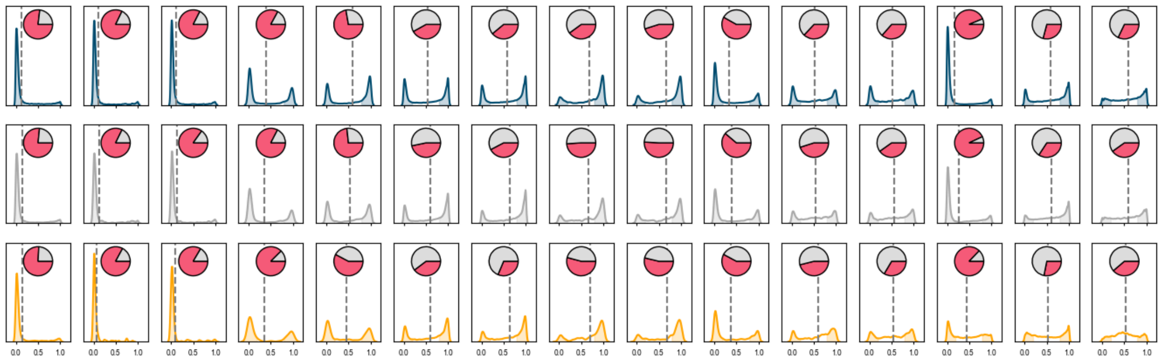

**Figure S9. pG4 density and pG4 enrichment *vs*. GC content.** The (A) pG4 density and (B) pG4 enrichment in human *versus* the GC content using 5-Mb windows. (C) And the models regressed over G4 enrichment and GC content for different window sizes. The N represents the total number of points present for each fit. (D) The distribution of residuals from the regression of G4 enrichment over GC content in 5-Mb windows.

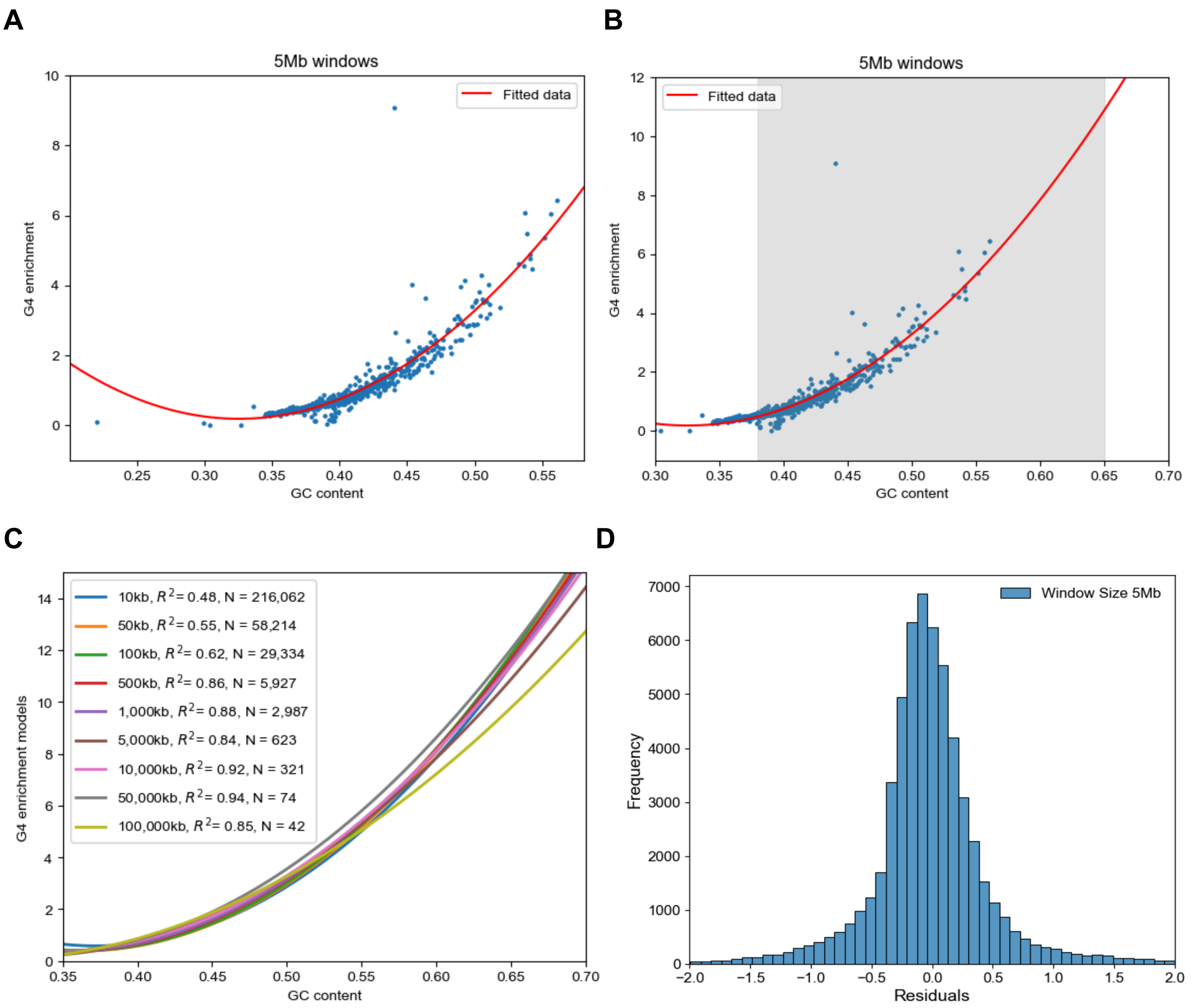

**Figure S10. Test for difference in the proportion of hypomethylated fraction of pG4s in different functional categories.** The heatmap for the difference in the proportion of hypomethylated fraction of pG4s in different functional categories between pG4 evolutionary groups at (A) bonobo, (B) chimpanzee, (C) human, (D) gorilla, (E) B. orangutan, and (F) S. orangutan. (ns: not significant, *: p-value<0.05, **: p-value<0.01, ***: p-value<0.001, two-tailed test of proportions).

**A**

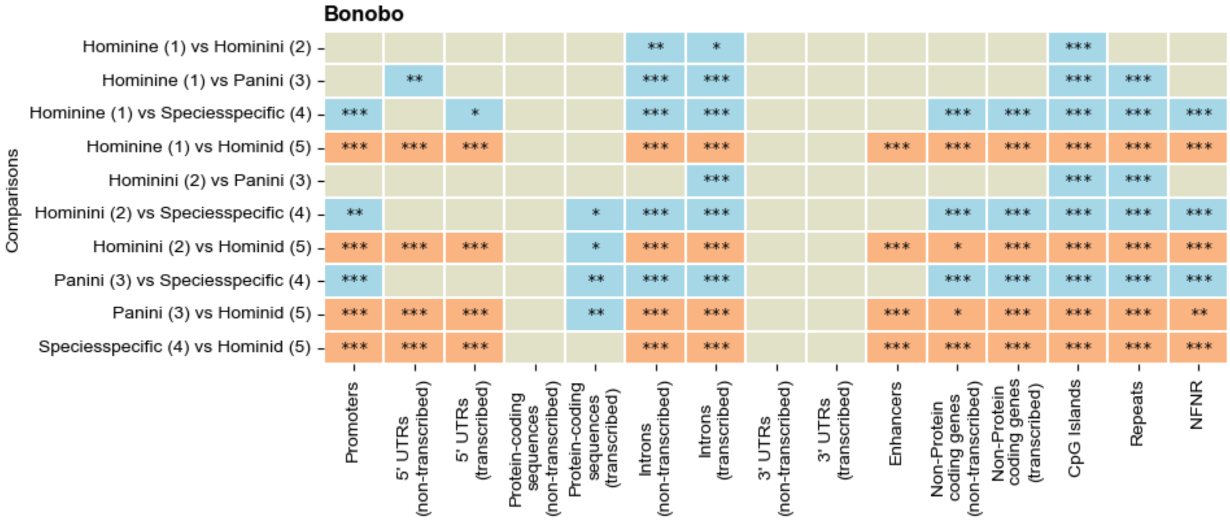


**B**

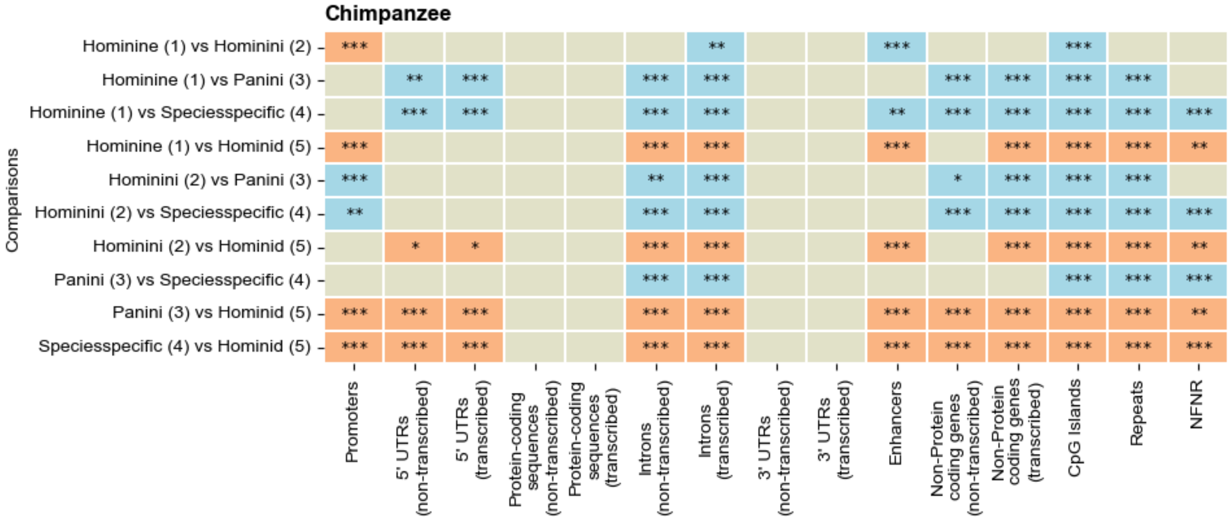

C

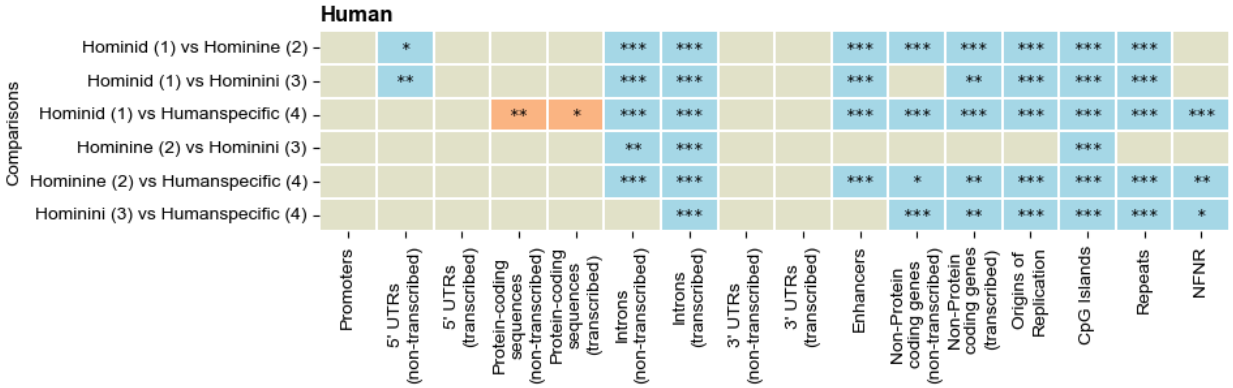
Human
Comparisons
Hominid (1) vs Hominine (2)
Hominid (1) vs Hominini (3)
Hominid (1) vs Humanspecific (4)
Hominine (2) vs Hominini (3)
Hominine (2) vs Humanspecific (4)
Hominini (3) vs Humanspecific (4)
Promoters
5' UTRs (non-transcribed)
5' UTRs (transcribed)
Protein-coding sequences (non-transcribed)
Protein-coding sequences (transcribed)
Introns (non-transcribed)
Introns (transcribed)
3' UTRs (non-transcribed)
3' UTRs (transcribed)
Enhancers
Non-Protein coding genes (non-transcribed)
Non-Protein coding genes (transcribed)
Origins of Replication
CpG Islands
Repeats
NFNR

D

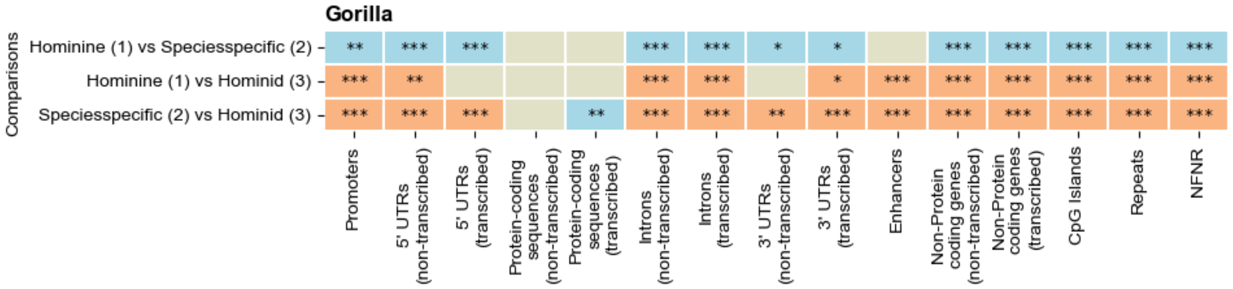
Gorilla
Comparisons
Hominine (1) vs Speciesspecific (2)
Hominine (1) vs Hominid (3)
Speciesspecific (2) vs Hominid (3)
Promoters
5' UTRs (non-transcribed)
5' UTRs (transcribed)
Protein-coding sequences (non-transcribed)
Protein-coding sequences (transcribed)
Introns (non-transcribed)
Introns (transcribed)
3' UTRs (non-transcribed)
3' UTRs (transcribed)
Enhancers
Non-Protein coding genes (non-transcribed)
Non-Protein coding genes (transcribed)
CpG Islands
Repeats
NFNR

E

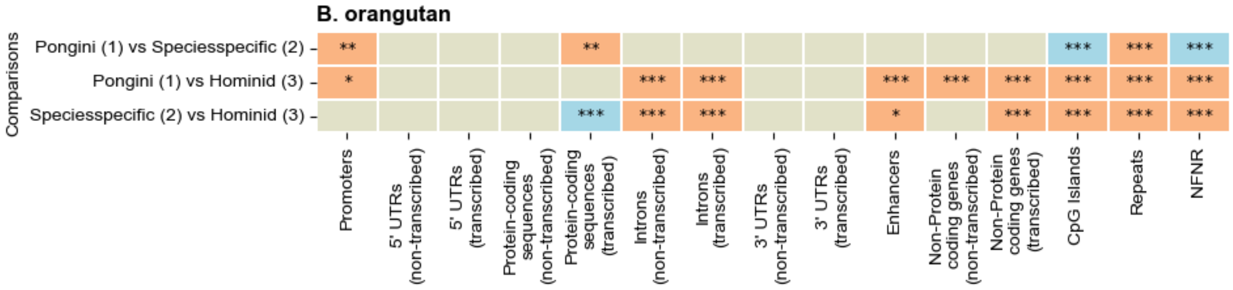
B. orangutan
Comparisons
Pongini (1) vs Speciesspecific (2)
Pongini (1) vs Hominid (3)
Speciesspecific (2) vs Hominid (3)
Promoters
5' UTRs (non-transcribed)
5' UTRs (transcribed)
Protein-coding sequences (non-transcribed)
Protein-coding sequences (transcribed)
Introns (non-transcribed)
Introns (transcribed)
3' UTRs (non-transcribed)
3' UTRs (transcribed)
Enhancers
Non-Protein coding genes (non-transcribed)
Non-Protein coding genes (transcribed)
CpG Islands
Repeats
NFNR

F

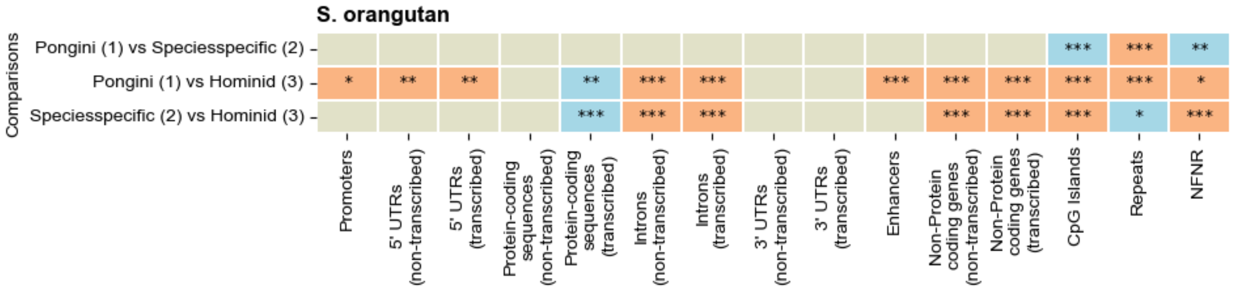
S. orangutan
Comparisons
Pongini (1) vs Speciesspecific (2)
Pongini (1) vs Hominid (3)
Speciesspecific (2) vs Hominid (3)
Promoters
5' UTRs (non-transcribed)
5' UTRs (transcribed)
Protein-coding sequences (non-transcribed)
Protein-coding sequences (transcribed)
Introns (non-transcribed)
Introns (transcribed)
3' UTRs (non-transcribed)
3' UTRs (transcribed)
Enhancers
Non-Protein coding genes (non-transcribed)
Non-Protein coding genes (transcribed)
CpG Islands
Repeats
NFNR

**Figure S11. pG4 methylation profiles of CHM13.** (A) The methylation profile for pG4s in the CHM13 cell line, and (B) their corresponding pairwise significance of hypomethylated G4 proportion. (C) The significance between the hypomethylated fraction for HG002 hominid evolutionary group versus the CHM13 hominid evolutionary group. (ns: not significant, *: p-value<0.05, **: p-value<0.01, ***: p-value<0.001, two-tailed test of proportions).

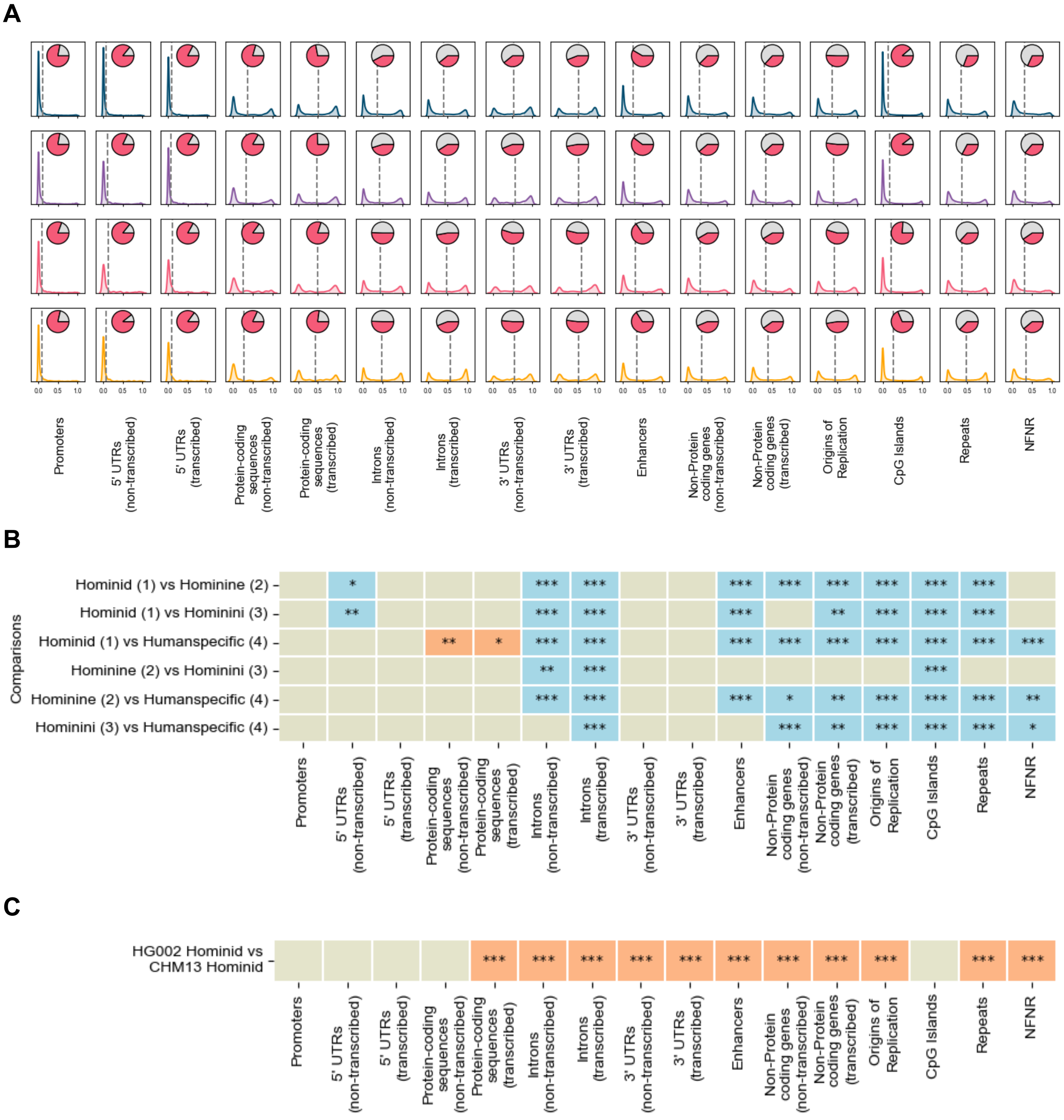

**Figure S12. Workflow of G4 discovery pipeline, pairwise genome alignments, and generation of connected graphs for shared and species-specific pG4s.** Great ape T2T genomes were used for pG4 prediction through the G4 discovery pipeline. Using the genomes, pairwise alignments were performed between the great ape chromosomes with human homology information. MAP-SEA was used to map the pG4s onto alignments and infer the profile of pG4 sharing. pG4s absent in alignment or present as a paired edge were considered as unaligned and aligned species-specific, respectively. The final pG4 database comprised both shared and species-specific pG4s.

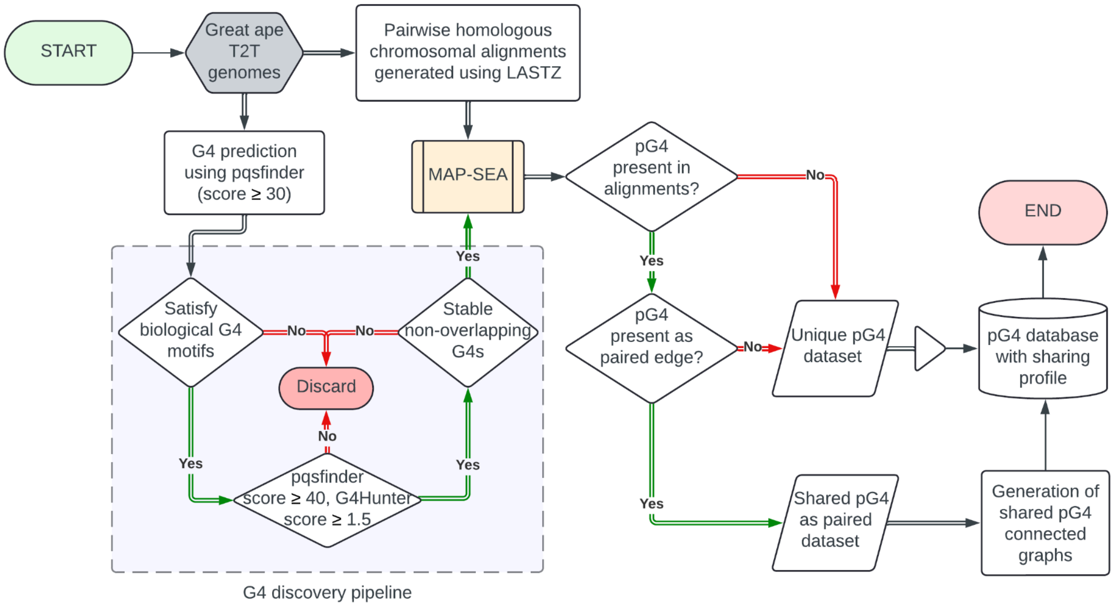
